# Neighborhood-statistics reveal complex dynamics of song acquisition in the zebra finch

**DOI:** 10.1101/595512

**Authors:** Sepp Kollmorgen, Richard Hahnloser, Valerio Mante

## Abstract

Motor behaviors are continually shaped by a variety of processes such as environmental influences, development, and learning^1,2^. The resulting behavioral changes are commonly quantified based on hand-picked features^3–10^ (e.g. syllable pitch^11^) and assuming discrete classes of behaviors (e.g. distinct syllables)^3–5,9,10,12–17^. Such methods may generalize poorly across behaviors and species and are necessarily biased. Here we present an account of behavioral change based on nearest-neighbor statistics^18–23^ that avoids such biases and apply it to song development in the juvenile zebra finch^3^. First, we introduce the concept of *repertoire dating*, whereby each syllable rendition is *dated* with a “pseudo” production-day corresponding to the day when similar renditions were typical in the behavioral repertoire. Differences in pseudo production-day across renditions isolate the components of vocal variability congruent with the long-term changes due to vocal learning and development. This variability is large, as about 10% of renditions have pseudo production-days falling more than 10 days into the future (*anticipations*) or into the past (*regressions*) relative to their actual production time. Second, we obtain a holistic, yet low-dimensional, description of vocal change in terms of a *behavioral trajectory*, which reproduces the pairwise similarities between renditions grouped by production time and pseudo production-day^24^. The behavioral trajectory reveals multiple, previously unrecognized components of behavioral change operating at distinct time-scales. These components interact differently across the behavioral repertoire—diurnal change in regressions undergoes only weak overnight consolidation^4,5^, whereas anticipations and typical renditions consolidate fully^2,6,25^. Our nearest-neighbor methods yield model-free descriptions of how behavior evolves relative to itself, rather than relative to a potentially arbitrary, experimenter-defined, goal^3–5,11^. Because of their generality, our methods appear well-suited to comparing learning across behaviors and species^1,26–32^, and between biological and artificial systems.

## RESULTS

Juvenile male zebra finches acquire complex, stereotyped, vocalizations through a months-long process of sensory-motor learning^3,28,33–38^. We obtained continuous audio recordings between 40 days post hatch (*dph*) and up to 120 dph (70.3±14.9 consecutive days, mean and STD across 7 birds). Birds were isolated from other males after birth and live-tutored between ages 44 and 67 (Fig. 1A, Methods). Audio recordings were band-passed (0.5k-8kHz) and segmented into individual syllable renditions represented as song spectrograms (Fig 1B, between 563124 and 1260715 renditions per bird). During development, both syllable order, i.e. syntax, as well as the spectral structure of individual syllables are altered^39^. Largely independent mechanisms, with distinct anatomical substrates, may underlie learning of these two properties of the song^40,41^. Here we focus on characterizing change in spectral structure.

**Figure 1.**
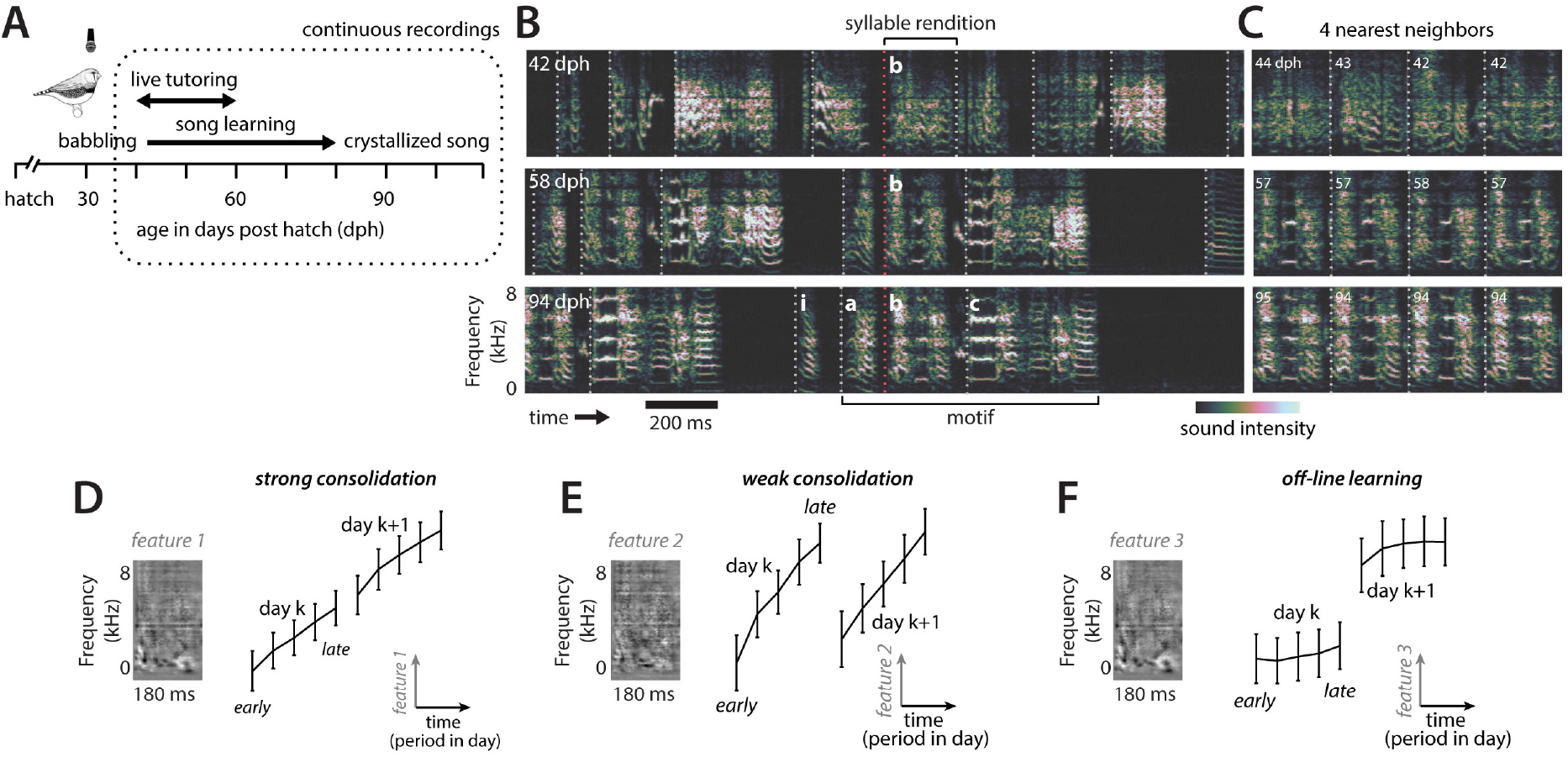
Vocal learning and motor variability in Zebra finches. (**A**) Vocal development in male Zebra finches. Tutoring by an adult male started between 39 and 47 days post hatch (dph) and lasted 10 to 20 days. (**B**) Example vocalizations of a single bird at three developmental stages (rows) represented as song spectrograms. Large changes in vocalizations are triggered by the exposure to tutor song. Song gradually transitions from largely unstructured subsong (top) to highly structured crystalized song (bottom). The onset of each individual syllable rendition was identified by segmenting vocalizations based on RMS-power threshold crossings (dotted lines mark segment onsets). As song becomes more structured, individual renditions fall into increasingly well-defined categories (syllables i, a, b, c; Extended Data Fig. 1) and are grouped into a stereotyped sequence of 3-8 syllables, called a motif, which is repeated up to several thousand times a day and resembles the tutor song (middle and bottom). Red dotted lines mark the onsets of example renditions of syllable b from the three developmental stages. (**C**) The four nearest-neighbors for the example rendition of syllable b in the corresponding row in (B). Nearest-neighbors were defined within the complete recordings for this bird based on Euclidian distance between spectrogram segments. Numbers indicate the production time (dph) of each nearest neighbor. Nearest-neighbors need not be produced on the same day as the corresponding example rendition in (B). (**D-F**) Time evolution of three spectral features of syllable b during days 57 and 58 dph. Each feature (left) corresponds to a linear kernel capturing change on a specific time-scale; the black curves (right) show the average projection of renditions of syllable b onto the corresponding feature (see Methods & Extended Data Fig. 8). These three features of the same syllable simultaneously undergo different patterns of overnight consolidation. (**D**) A feature reflecting the long-term developmental trend. (**E**) A feature reflecting within-day change. (**F**) A feature reflecting across-day change.

### Learning curves differ across behavioral features

In the context of learning, behavioral change is often described through “learning curves”, which depict how particular features of the behavior, like overall task-performance or syllable pitch, change over time^3–6,42^. Learning curves can provide a parsimonious description of change for behaviors that are intrinsically low-dimensional, like reflexive eye-movements^43^, or that are constrained to lowdimensional output spaces, like the position of a planar manipulandum^1,2,7,8,29^. Change in complex, high-dimensional behaviors like singing^3^, however, can be difficult to capture with “learning-curves” alone.

To illustrate this point, we compute the time evolution of a few hand-picked spectral features of a single birdsong syllable at a particular developmental stage. Here we consider three linear features (Fig. 1D-F, insets; see methods for the definition of the features). The strength of each feature for a given rendition corresponds to the dot product of the rendition spectrogram with the linear kernel defining the feature. The corresponding “learning curve” is obtained by plotting feature strength over time, across two adjacent days (Fig. 1D-F, black lines).

The three chosen features result in substantially different descriptions of the time-course of change for the same syllable. The first feature becomes progressively stronger during the day, and these changes are consolidated overnight (Fig. 1D; strong consolidation). The second feature also becomes stronger during the day, but these changes are largely lost overnight (Fig. 1E; weak consolidation). The third feature does not change much during the day but strengthens overnight (Fig. 1F; offline learning). Notably, strong consolidation^6,25^, weak consolidation^4,5^, and offline learning^6,44,45^ have all been reported in the past^30^, albeit in different behaviors and species. Figures 1D-F show that these different patterns of change can coexist within a single complex behavior.

Past studies relying on only one or a few hand-picked features of behavior, like pitch^3^ or entropy variance^4,5^, may thus have provided a biased, and possibly misleading, picture of the time course of change during vocal development. One approach to avoiding this bias might be to attempt to identify all features undergoing change. This would be cumbersome, since the relevant features are likely to differ both across syllables and stages of development^3^. Because the relevant features differ across syllables, the song would additionally have to be clustered into distinct syllables. This requirement is particularly problematic for behavior that does not appear to be clustered (e.g. juvenile song) and when the number of clusters changes over time (Extended Data Fig. 1). A related approach is to directly track the similarity of vocalizations to the tutor song^11^. However, the “goal” of vocal development reflects not just the tutor song, but also innate song priors of largely unknown nature^46^. Focusing on similarity to tutor song thus also leads to a biased description of change, as it potentially blends out critical components of the learning process.

### Nearest-neighbor based non-parametric assessment of behavioral change

Here we develop a novel, non-parametric, *high-dimensional* characterization of behavioral change that does not rely on features, but rather provides a “holistic” description of behavioral change based on nearest-neighbor statistics^19–21,47,48^. We analyze spectrogram segments of fixed duration (e.g. 60ms) aligned to syllable onset. The segments are represented as real valued vectors *x_i_* where *i* indexes all renditions produced by a bird. The nearest-neighbor of syllable rendition *x_i_* is defined as the data point. that is closest to it based on Euclidean distance in spectrogram space: *NN_i_* = *argmin_j_* |*x_i_ – x_j_*|^2^, where *i* and *j* range over the entire dataset (Fig. 1C). The set of *k* nearest-neighbors of rendition *i* defines its *local neighborhood.* Each data point has an associated *production time, t_i_* ∈ ℝ, i.e. the bird’s age when singing *x_i_*. Larger values of *k* result in more global views of the learning process. However, *k* should be small enough to not mix different clusters within a neighborhood (Extended Data Fig. 1C).

**Figure 2.**
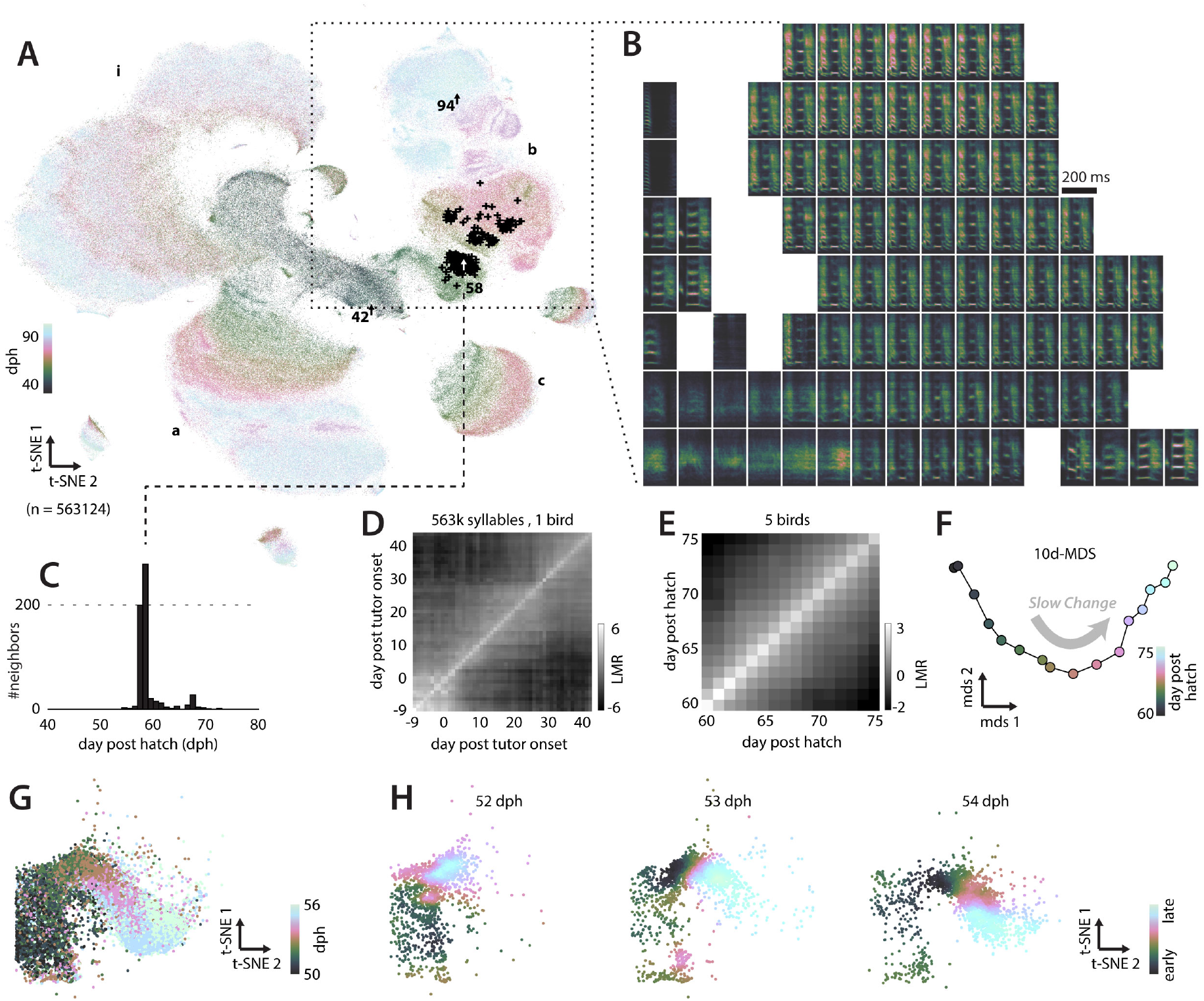
Change in vocalizations on multiple time-scales. (**A**) t-SNE embedding of the continuous behavioral recording for the example bird in Fig. 1. Each data point corresponds to a syllable rendition. Late in development, data points fall into separate clusters (syllables i, a, b, and c; labels). The three example renditions of syllable b from Fig. 1B are indicated with arrows. Crosses mark the 600 nearest-neighbors of the example rendition from day 58. (**B**) Average spectrograms for different locations in the t-SNE embedding, obtained by averaging all spectrograms associated with data points at the corresponding grid location in (A). Points at nearby locations in (A) have similar spectrograms. (**C**) Histogram of production times over the 600 nearest-neighbors of the example syllable in (A) (black crosses). (**D**) Mixing-matrix based on all data points in (A). Data points are subdivided into days. Each column of the matrix represents a histogram of production times (as in C), averaged over all neighborhoods of points with a day (horizontal axis) and normalized with respect to a shuffling null-hypothesis (LMR: logarithm of mixing ratio, in base 2). Here production time is given relative to tutor onset. (**E**) Average mixing-matrix over 5 birds for days 60-75 dph. (**F**) Reconstructed behavioral trajectory based on (E), inferred with 10-d multi-dimensional scaling. Each point corresponds to a day. Only the 2 dimensions capturing most variance in the trajectory are shown (Extended Data Fig. 4). (**G**) Across-day change in vocalizations. Magnified cutout from (A) (bottom-left region of the dashed square). Coloring differs from (A); only points from days 50-56 are shown. (**H**) Within-day change in vocalizations. Points from (G) are shown separately for three individual days (top labels) and colored based on production time within the day (early to late; color bar). Vocalizations change within each day—early vocalizations (dark green) are more similar to vocalizations from previous days (dark green points in A); late vocalizations (light blue) are more similar to vocalization from future days (light blue in A).

To explore visually how spectrograms change with learning, we first represent the data for a given bird with Barnes Hut t-stochastic neighbor embedding^23^. This non-linear dimensionality reduction technique visualizes high dimensional data by finding a low dimensional arrangement of the data points that predominantly preserves local neighborhoods. Each point in such a low dimensional *embedding* (Fig. 2A) corresponds to a spectrogram segment *x_i_* and different locations in the embedding space correspond to different vocalization types (Fig. 2B). The embedding suggests that vocalizations change from undifferentiated subsong^49^ (Fig. 2A, middle) to clearly differentiated syllables falling into at least four categories (Fig. 2A, syllables *a, b, c* and introductory note *i*; same labels as in Fig. 1B). Initially, individual syllable renditions are highly variable, and averages of neighboring syllables are undefined and blurry (Fig. 2B, bottom-left panels) but over time stereotyped syllable types emerge (Fig 2B, e.g. top row). This emergence of clustered syllables from un-clustered subsong is confirmed by standard clustering algorithms (Extended Data Fig. 1). Note that the embedding does not reproduce all local structure in the underlying data, as nearest neighbors in the embedding space are not necessarily nearest neighbors in the full high dimensional data (Fig. 2A, black crosses mark high-d neighbors). Behavioral change should therefore not be quantified in lowdimensional embeddings, but rather directly on the high-dimensional data.

Specifically, we quantify behavioral change by analyzing the composition of local neighborhoods^18,20,47^ (Extended Data Fig. 2). For each data point, we compute the histogram of production times over its neighborhood (the “neighborhood production times”; Fig. 2C) and summarize all histograms as a *neighborhood mixing-matrix* (Fig 2D; Extended Data Fig. 2C). Each value of this matrix represents the similarity between vocalizations from two production times. Deviations from zero indicate that vocalizations from the corresponding production times are more (> 0) or less (< 0) similar (i.e. mixed at the level of neighborhoods) than expected from a shuffling null-hypothesis.

The mixing-matrix for the example bird depicts an initial, largely stationary subsong phase (Fig. 2D, bottom-left; all pairs of days<0 have high similarity). Marked changes occur at tutor onset (Fig. 2D, days=0) and more gradual changes afterwards (Fig. 2D, days>0; similarity largest along the diagonal). A similar progression is observed across all recorded birds (Extended Data Fig. 3).

To obtain a more accessible representation of the mixing matrix, we use non-metric multi-dimensional scaling^50^ to represent the similarity structure between vocalizations from different production times as a *behavioral trajectory* (Fig. 2F; see Methods). Each point on the inferred 10-dimensional trajectory represents all vocalizations produced on a given day and the pairwise distances between points faithfully represent the measured similarity structure (Extended Data Fig. 4A). Here we focus on a 16-day-phase of gradual change occurring midway through development (Fig. 2E). During this phase, the behavioral trajectory is structured differently at fast and slow timescales (Extended Data Fig. 4). The 2d-projection of the trajectory that explains maximal variance emphasizes the slow components of behavioral change (Fig. 2F).

These slow components of change reflect the long-term changes in behavior that are typically equated to learning and development. By contrast, fast components may reflect metabolic, neural, or other changes within single days that are not congruent with the slow changes resulting from learning or development. In the following we abstractly refer to the slow components as the *direction of slow change* (*DSC*).

The behavioral trajectory (Fig. 2F) has three notable properties. First, it provides a characterization of change that does not require an explicit clustering step to categorize vocalizations into syllables. Movement along the behavioral trajectory captures both the progressive differentiation of vocalizations into distinct syllables and continuous changes in the structure of individual syllables (Fig. 1, Fig. 2A, B; Extended Data Fig. 1). Second, movement along the trajectory potentially reflects simultaneous change in many spectral features, which need not be identified explicitly. Third, behavioral change is defined by comparing the juvenile song to itself across different developmental stages, rather than to a tutor song. Thus the behavioral trajectory also reflects innate song priors that can result in crystallized song deviating from the tutor song^46^, as well as additional change due to other developmental processes.

### Repertoire dating reveals repertoire extent and consolidation

The slow components of change in Fig. 2F do not account for all the dynamics in the behavior. The lowdimensional t-SNE embedding reveals that renditions produced on a given day are highly variable and that behaviors from nearby days overlap considerably (Fig. 2H). This variability suggests that individual renditions from a day are widely spread along the behavioral trajectory. Underlying this variability is a clear temporal dynamic. Behavior changes systematically over the course of hours (Fig 2H) and these within-day changes appear to partly mimic the gradual change across days (Fig. 2G).

Critically, the direction of slow change (DSC) provides a principled means to parameterize behavior by its “maturity”, which in turn can be used to relate changes occurring at different timescales. Particularly mature, or *anticipatory*, behaviors resemble renditions mostly produced in the future. *Anticipations* can be thought of as lying further along the trajectory than regressive behaviors, which instead resemble renditions produced in the past. The position along the DSC for a given rendition can be determined by means of the neighborhood production times (Fig. 2C, Fig. 3A). Anticipations and regressions predominantly have neighbors that were produced in the future or in the past, whereas renditions that were *typical* for a given developmental stage mostly have neighbors produced on the same or nearby days (examples in Fig. 3A, all produced on day 70). The median of the neighborhood production times defines the “*pseudo” production-day* (*pPD*) for each rendition. The pseudo production-day can serve to place each rendition along the DSC (Fig. 3C, horizontal axis; Fig. 3D), i.e. to *date* it with respect to the progression of development. For typical renditions, the pseudo production day matches the true production time (Fig. 3C,D; pPD=70), whereas for anticipations it is larger (pPD>70) and for regressions smaller (pPD<70). We refer to this parameterization of behavior as “*repertoire dating*” (in analogy to methods like radiocarbon-*dating*).

**Figure 3.**
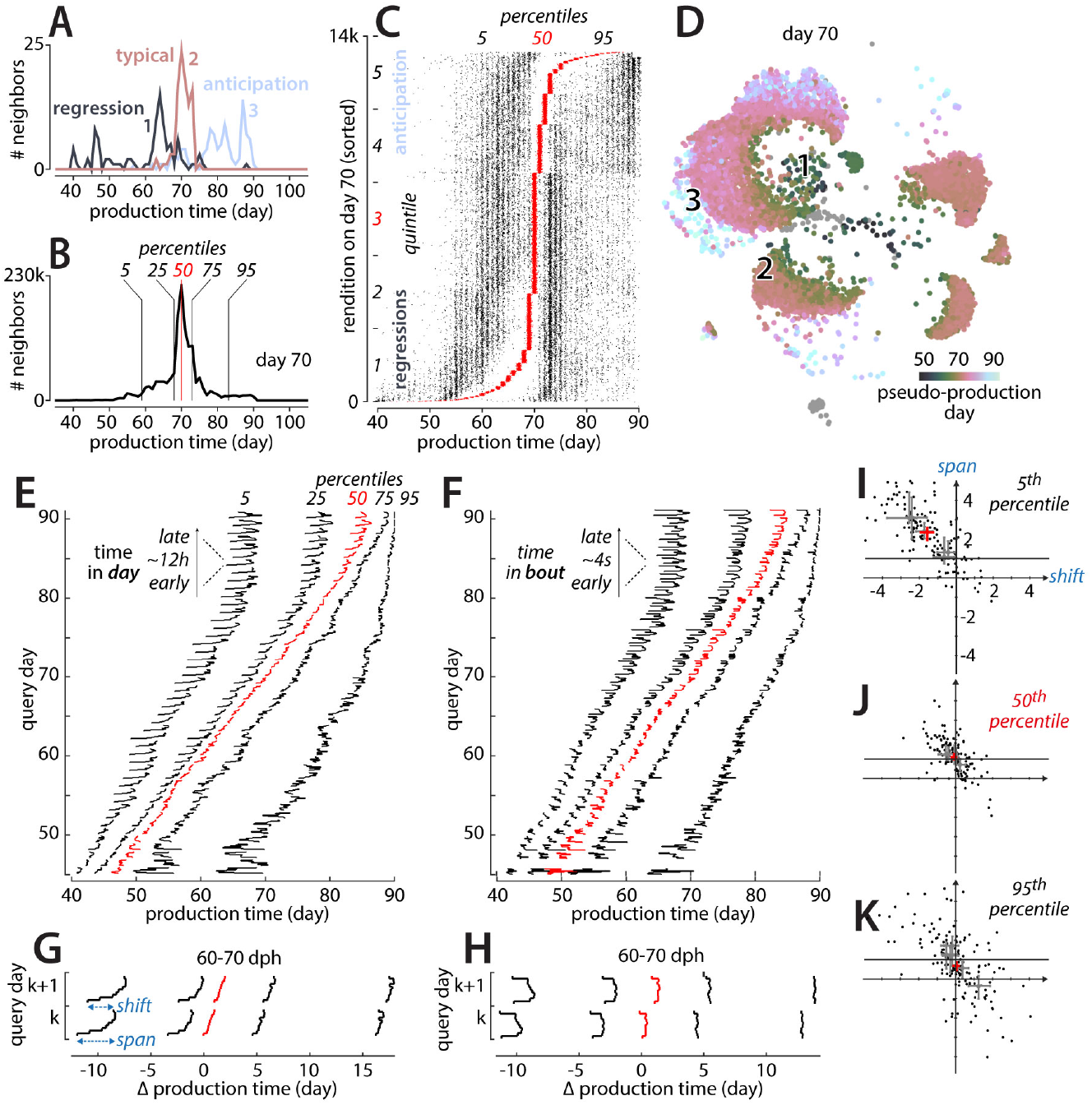
Variability of individual renditions within a dynamic behavioral repertoire. (**A**) Neighborhood production-time histograms (over 600 nearest-neighbors) for three renditions from day 70 (analogous to Fig. 2C). Rendition 2 is typical for day 70 (most neighbors lie in the same or adjoining days); renditions 1 and 3 are a “regression” and an “anticipation” (typical of days in the past or future, respectively). Data in A-D from the same bird as in Fig. 2. (**B**) Pooled neighborhood productions times for day 70 (sum of all histograms as in (A) for day 70). Percentiles (vertical lines) quantify the extent of the behavioral repertoire of day 70 along the direction of slow change (compare Fig. 2F). (**C**) Percentiles (5^th^, 50^th^, 95^th^) of neighborhood production-time histograms (as in A) for day 70. Each row is a rendition. Rows are sorted by the 50^th^ percentile, i.e. the pseudo-production day (pPD). The left, middle (red), and right dots mark the corresponding percentiles. A small random horizontal shift was added to each dot for visualization. (**D**) All renditions of day 70 (subset of points from Fig. 2A). Color corresponds to the pPD (50^th^ percentile in C). Anticipations (pPD>70) and regressions (pPD<70) occur at locations corresponding to vocalizations typical of later and earlier developmental stages (compare to Fig. 2A). (**E**) Within and across-day changes in behavioral repertoire along the direction of slow change. The behavioral repertoire is computed separately for 10 consecutive periods in any day (arrow in inset). Each row shows percentiles of the pooled neighborhood production times (B) for a given “query” day and period. Average over 5 birds. (**F**) Within-bout changes in behavioral repertoire. Analogous to (E), but renditions from any query day are binned into 10 periods depending on time of occurrence within the corresponding singing bout (arrow in inset). (**G**) Average within and across-day changes (average over days 60-70 in E). Production time is expressed relative to the query day. (**H**) Average within-bout changes (average over days 60-70 in F). Analogous to (G). (**I**) For each day we define a within-day span (first to last period of day k) and an overnight shift (first period of day k+1 to last period of day k; blue arrow in G) for the 5^th^ percentile (regressions) for given day (dph 50-80) and bird; data from 5 birds. Gray crosses and the red cross indicate median and bootstrap confidence intervals for each bird and across all birds, respectively. Span is large and positive, shift is large and negative, indicating week consolidation. (**J**) Analogous to (I), but for the median (typical renditions). Shift is close to zero, and span close to 1, indicating strong consolidation. (**K**) Analogous to (I), but for the 95^th^ percentile (visions).

The distribution of pseudo production-days across all renditions in a day (Fig. 3C) provides an estimate of the extent of motor variability along the DSC. By his measure, variability is large—the most extreme regressions are backdated more than 10 days into the past, and the most extreme anticipations are postdated more than 10 days into the future. The extent of motor variability along the DSC is also reflected in the *pooled* production times (Fig. 3B), obtained by combining the neighborhood production times of all renditions from a given period. For day 70 of the example bird, the histogram of pooled production times peaks around day 70, reflecting that most renditions are typical for the developmental stage when they were produced (Fig. 3B, 50^th^ percentile at or close to 70). However, the tails of the histogram extend far into the future and into the past (Fig. 3B), reflecting the substantial fraction of anticipations and regressions in the behavioral repertoire.

We use the pooled production times to quantify changes in the behavior at the time scale of hours. We subdivide each day into 10 consecutive periods and compute pooled production times separately for each period. The evolution of behavior within and across days can then be assessed for the entire duration of development by tracking the percentiles of the pooled production times over time (Fig. 3E). The evolution of each percentile is akin to a “learning curve” (e.g. Fig. 1D-F) for a specific part of the behavioral repertoire (e.g. typical renditions, 50^th^ percentile; extreme anticipations, 95^th^ percentile). The curve for each percentile captures the progress along the DSC (Fig. 3E, horizontal axis) as a function of time (Fig. 3E, vertical axis). We validated this characterization of behavioral change on simulated behavior mimicking vocal development in zebra finches (Extended Data Fig. 5).

The percentile curves reveal a strikingly different time-course of change for anticipations, regressions, and typical renditions. Typical renditions move gradually along the DSC throughout the day (Fig. 3E, G, red; horizontal axis). Changes acquired during the day are, on average, *fully consolidated* overnight, i.e. typical renditions produced early on day k+1 fall onto a similar location along the DSC as typical renditions produced late on day k (Fig. 3G, red; Fig. 3J; compare to Fig. 1D). A similar or smaller degree of within-day-change is observed for anticipations (Fig. 3E, G; 75^th^ and 95^th^ percentiles). The dynamic of consolidation for regressions is very different (Fig. 3E, G; 5^th^ and 25^th^ percentiles). Within each day, regressions move by a large distance, but this change is only weakly consolidated overnight (as in Fig. 1E; Fig. 3G; 5^th^ and 25^th^ percentiles; Fig. 3I). This implies that the worst renditions improve markedly throughout a day, more so than typical renditions or anticipations, but these improvements are mostly lost overnight.

Notably, movement along the DSC also occurs at much faster timescales than hours, namely within bouts of singing. We define a bout as a group of vocalizations preceded and followed by extended periods of non-singing. We require pauses between adjacent bouts of at least 2.5s, resulting in an average bout duration of 3.81±0.83s (see Methods). On average across birds, 99% of all renditions from a day occur within bouts. We identify within-bout changes (Fig. 3F,H) in the same way as within-day changes (Fig. 3E,G)—we subdivide each bout into 10 consecutive periods, compute pooled production times for each period (over all bouts in a day), and track change through the corresponding percentiles. Within bouts, large, consistent changes along the DSC occur at the regressive tail of the behavioral repertoire—vocalizations are most regressive at the onset of a bout, become substantially less regressive as the bout progresses, only to become more regressive again before the end of the bout (Fig. 3F, H; 5^th^ percentile). Similar, albeit weaker, changes occur for typical renditions (Fig. 3F, H; red).

The same changes in song maturity can be observed when short bouts (duration 2.30±0.54s) or long bouts (duration 6.28±1.73s) are considered separately (Extended Data Fig. 6A, B), suggesting that the end of a bout predictably follows the observed decrease in song maturity. Importantly, these within-bout changes reflect systematic changes in the structure of the underlying spectrograms, rather than changes in the overall frequency of production of different syllables across bout periods (Extended Data Fig. 6C, D). Birds may thus actively monitor their singing behavior^51^ and stop singing when the quality of their vocalizations decreases.

### Stratified neighborhood mixing matrices reveal misaligned behavioral components

The pseudo production-day, by construction, can only reveal within-day and within-bout changes that recapitulate, on a faster timescale, changes that are also apparent at the resolution of days. For example, consider a hypothetical syllable that undergoes change with respect to a first feature that changes within days, from morning to evening, but not over days (as in Fig. 1E); and a second feature that changes over days, but not within days (as in Fig. 1F). The pseudo production-day would, on average, be identical for morning and evening renditions, and thus not reflect the existence of within-day change along the first feature. We refer to components of change that are reflected in the pseudo production-day as being *aligned* with the direction of slow change (DSC) and components that are not reflected in it as being *misaligned* or orthogonal.

To shed light on the relationship and size of aligned and misaligned components, we construct a representation of the data that can resolve both components of change. We define a *stratified* mixing matrix, which combines the concept of a neighborhood mixing-matrix (e.g. Fig. 2D, E) with repertoire dating (Fig. 3). Specifically, we use repertoire dating to subdivide each day’s behavioral repertoire into 5 *strata*, by assigning each rendition to 1 of 5 quintiles based on its pseudo production-day (Fig. 3C, quintiles). Additionally, we also assign each rendition into 1 of 5 consecutive production periods, analogous to the 10 periods used in Fig. 3E-H. We then place the renditions into 1 of 25 bins, defined by all pairwise combinations of strata and production periods (the binning is nested, Fig. 4). The full stratified mixing matrix measures similarity between the 50 bins obtained by combining data from two adjacent days (Fig. 4A, B). We average these stratified mixing matrices across pairs of days and birds.

**Figure 4.**
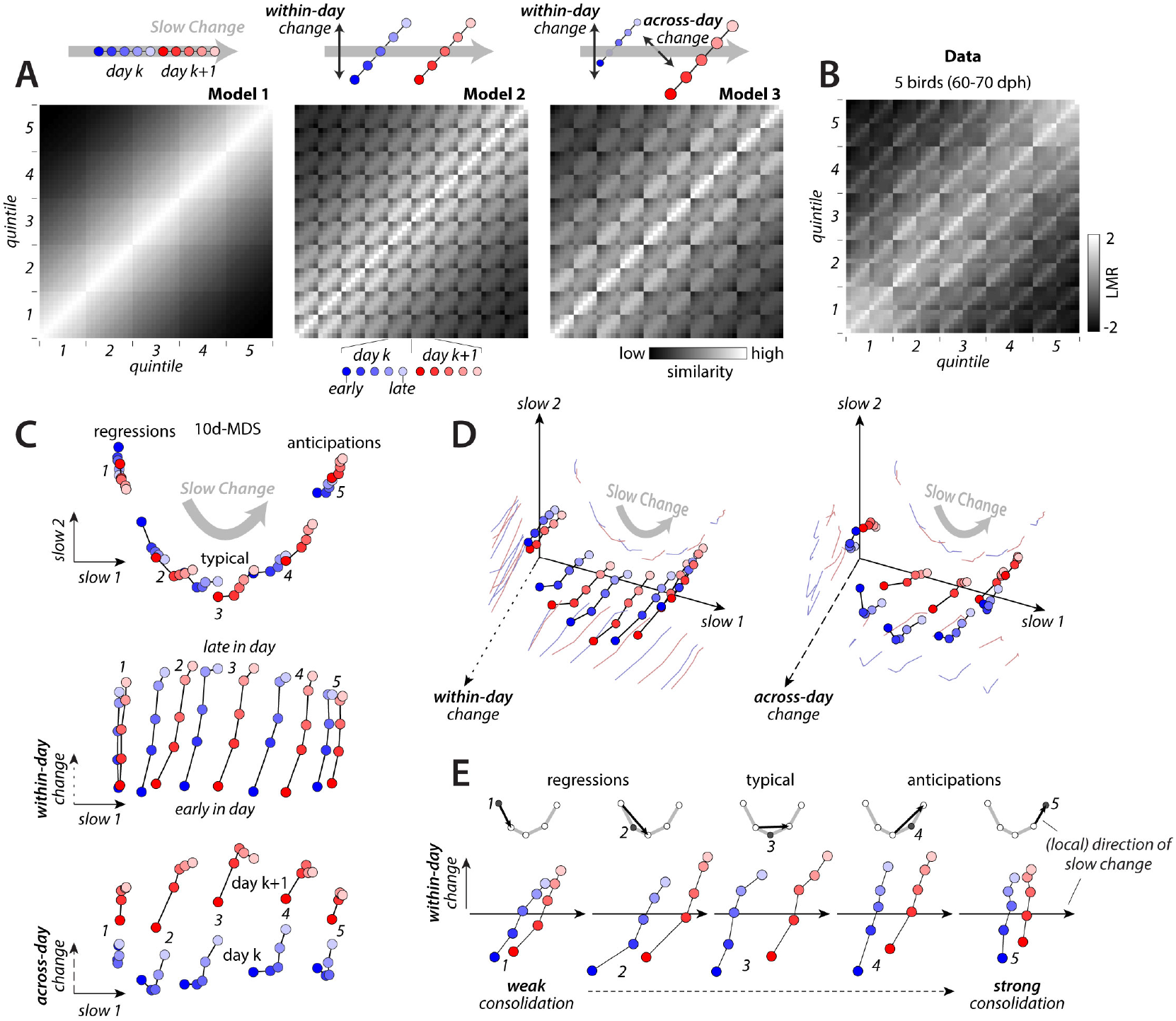
Multiple components of behavioral change during sensory-motor learning. Stratified mixing matrices and corresponding behavioral trajectories. Renditions are assigned to 1 of 5 strata based on pseudo-production day (quintiles 1-5, as in Fig. 3C) and into 5 periods based on their production time in a day (early to late). (**A**) Simulated stratified mixing matrices for three models corresponding to different alignments of within-day and across-day change with the direction of slow change (DSC) (schematic behavioral trajectories on top). (**B**) Measured stratified mixing matrix, averaged over day-pairs within the range 60-70 dph and across birds. (**C**) Stratified behavioral trajectories obtained with 10-dimensional multi-dimensional scaling based on the mixing matrix in (B). Three orthogonal 2-d projections of the trajectories reveal the DSC (top), as well as the components of within-day change (middle) and across-day change (bottom) that are not aligned with the DSC (along directions of within-day change and across-day change, respectively; arrows). Labels 1-5 mark the five strata (1,2 = regressions; 3 = typical; 4,5 = anticipations). The full 10-d trajectories faithfully reproduce the similarity structure in the mixing matrix (MDS stress = 0.016). The 4-d subspace spanned by these three orthogonal 2-d projections captures 81% of the variance of the full trajectory. (**D**) Two additional 3-d projections of the trajectories in (C), showing combinations of both dimensions of slow change (C, top) with the direction of within-day change (left) or the direction of across-day change (right). (**E**) Overnight consolidation of within-day change across the DSC. Because the DSC varies across the behavioral repertoire and different strata lay at different locations (C, top; labels 1-5), the exact alignment of within-day change and the DSC is assessed best when computing separate projections for each stratum onto the local DSC. This local direction is defined as the vector spanning the average location of points from two appropriate strata (inset on top, black arrows; each point in the inset represents one stratum, same projection as in C). Consolidation along the local DSC is weakest for regressions and strongest for anticipations.

We compare the average stratified mixing matrix (Fig. 4B) with simulations that differ with respect to how change within a day and change across adjacent days are aligned with the DSC (Fig. 4A, top; Extended Data Fig. 7). In simulated model 1, change only occurs along the DSC, meaning that development can be thought of as a “1-dimensional” process. In model 2, change within a day involves both a component along the DSC, as well as a consistent component that is not aligned to it (Fig. 4A, within-day change). Model 3 includes an additional misaligned component of change compared to model 2. Adjacent days are separated not only along the DSC, but also along a direction orthogonal to it (Fig. 4A, Model 3, across-day change). For simplicity, here we do not simulate distinct consolidation patterns across the behavioral repertoire, but rather let all strata undergo full overnight consolidation. The prominent “stripes” along every other diagonal in the measured mixing matrix (Fig. 4B) indicate larger similarity between renditions from the same day than between renditions from adjacent days. This structure appears most consistent with model 3, suggesting that several misaligned components contribute to change at fast timescales.

We visualize these components through multi-dimensional scaling on the stratified mixing matrix by concurrently inferring 10-dimensional behavioral trajectories for all strata of the behavior. These inferred trajectories span more than two dimensions. To appreciate their structure, we thus consider several distinct 2-dimensional-projections of the inferred trajectories (Fig. 4C-E; labels 1-5 mark the different strata; Extended Data Fig. 7). The projection capturing most of the variance due to the 5 strata resembles Fig. 2F and reflects the component of slow change (Fig. 4C, top). Consistent with Fig. 3E, the change in behavior between adjacent days along the DSC (Fig. 4C, top; blue vs red points for the same stratum) is small compared to the spread of one day’s behavior along the same direction (e.g. blue points, strata 1-5). For each stratum, however, much of the change occurring within a day is misaligned with the DSC (Fig. 4C, middle; early vs. late separated along orthogonal dimension of within-day change). Yet another misaligned component is necessary to appropriately capture change across adjacent days (Fig. 4C, bottom, day k vs. k+1 separated along orthogonal direction of across-day change).

The trajectories also reflect the heterogeneous dynamics of consolidation across the behavioral repertoire (Fig. 4E). Because the behavioral trajectory is curved, the consolidation dynamic of each stratum is more faithfully assessed through a separate projection onto the “local” DSC (Fig. 4E, inset; black arrows). The 2d-projections spanned by this local direction and the direction of within-day change confirm the findings of Fig. 3G—for regressions, most of the gain along the DSC experienced during a day is lost overnight (Fig. 4E; strata 1&2; weak consolidation, Fig. 1E); for typical renditions (stratum 3) and anticipations (strata 4-5) the within-day gain instead is maintained or possibly increased upon, corresponding to strong consolidation or even “off-line” learning (Fig. 1D, F). Notably, here the distinct consolidation patterns are revealed for each stratum separately, whereas the analysis in Fig. 3G is based on an aggregate description of the behavioral repertoire (the pooled neighborhood production times, e.g. Fig. 3B).

The interplay of the identified fast and slow components of change explains the potentially puzzling finding that different features of a single syllable can undergo different patterns of change (Fig. 1D-F): the time evolution for a given feature is determined by the feature’s relative alignment with the DSC and the within-day and across-day change. That alignment in turn can differ from feature to feature. Indeed, the properties of aligned and misaligned components of change inferred from nearest-neighbor statistics can be replicated with a linear analysis^52,53^ based on spectral features chosen to capture change at specific timescales (Extended Data Fig. 8, 9). The pattern of overnight consolidation for a given feature, in particular, mainly reflects its degree of alignment with the DSC. Consolidation can be strong (complete alignment), weak (partial alignment), or reveal “offline learning” (partial alignment with the DSC but orthogonal to within-day change; Extended Data Fig. 8).

## DISCUSSION

Our finding that within-day changes are fully consolidated overnight across much of the behavioral repertoire is in line with studies of skilled motor learning in humans^31,44,45,54^ and of motor adaptation in humans^2,29^ and birds^6^, but appears at odds with past accounts of vocal learning in Zebra finches, which reported weak consolidation^4,5^. The latter results can be reconciled with our findings if one assumes that consolidation was assessed on a song feature (“song complexity”, measured as entropy variance^55^) that happens to be more aligned with the component of within-day change than with the direction of slow change (DSC) (like the feature in Fig. 1E; Extended Data Fig. 8). Notably, an increase in “song complexity” has also been observed during directed singing, as opposed to the undirected singing we study here^56^. This increase need not necessarily imply that juveniles can dramatically improve the maturity of singing during phases of exploitation compared to exploration, as previously proposed^56^. The difference in directed vs. undirected song may occur along the direction of within-day change, which would result in little gain along the DSC.

The identified components of behavioral change could reflect both neural and non-neural processes. Neural bias signals from area LMAN may contribute to the observed within-day change (as in “pitch-shifting” experiments^6,57^) whereas across-day change may reflect synaptic and circuit-level modifications occurring overnight^6^. Non-neural processes like circadian rhythms^5,58^, fatigue, or varying environmental factors could also contribute to the observed components of change. The strength of within-day and across-day change is strongest after tutor onset, and declines as the song becomes more mature (Extended Data Fig. 10), suggesting that they do play a role in learning. The component of slow change may also reflect developmental processes related merely to the growth of juvenile birds. A large contribution of growth-related or similar, non-specific, processes to this component would have to be reconciled with the observed stark differences in consolidation across the repertoire; an overnight reset along the DSC for regressions; and an interruption of change during the night in typical renditions (Fig. 3G, Fig. 3I-K, and Fig. 4E). Furthermore, movement along the DSC does not only occur on the slow timescales to be expected from developmental processes, but also on the much faster timescale of bouts (Fig. 3F, H). Nonetheless, behavioral analyses alone can ultimately not distinguish between neural and non-neural contributions to change.

Lacking a formal understanding of how species-specific priors and tutor song are combined to set the goals of learning, any characterization of the progress of learning, whether based on features or nearest neighbors, is not immune to contamination by developmental and other processes. Yet, an approach based on a few handpicked features is, a priori, necessarily more biased with respect to the entirety of behavioral change than our “intrinsic” characterization of learning as the direction towards which the song is moving. Hence, the fine-grained structure of behavioral change revealed here provides a novel, more precise and more general foundation for both theoretical^59–61^ and experimental studies into the neural processes of learning.

Our approach, based on nearest-neighbor statistics^19–22,47,48^ is largely complementary to approaches that rely on clustering behavior into distinct categories^10,12,15–17,62^. Whereas such model based, or parametric, methods fare better when an appropriate model is known^22^ the approach presented here is model-free and can be applied when no appropriate models are known. To date, the latter seems to be the case for many complex behaviors, including birdsong. Nearest-neighbors respect potential clustering structure in the data “per construction”, since they will typically lie within the same cluster (Extended Data Fig. 1). Foregoing an explicit clustering of the data can be advantageous since assuming the existence of clusters can constitute an unwarranted approximation^17,63,64^ and impede the characterization of behavior that appears not clustered, like juvenile Zebra finch song (Extended Data Fig. 1). Furthermore, determining precise cluster counts and precise cluster boundaries is an ill-defined problem even for data that appears well clustered^65,66^. Importantly, the definition of nearest-neighbors requires only a locally meaningful distance metric^67^, a much weaker requirement than a globally valid distance metric or the existence of a low dimensional feature space that globally maps behavioral space^68–72^. Because of these properties, the presented framework is very general, and applicable to almost any behavior, as well as other high dimensional data characterized by “labels” other than production time. The framework may thus provide an account of learning amenable to comparisons between different behaviors and model systems, including different species^26,73^ and artificial learning systems^74^.

## FUNDING

This research was supported by funding from the Simons Foundation (awards 328189 and 543013 to VM) and the Swiss National Science Foundation (award PP00P3_157539 to VM; award 31003A_182638 to RH).

## ETHICS OVERSIGHT

All experimental procedures were approved by the Veterinary Office of the Canton of Zurich.

## METHODS

### Vocalization Recordings

Vocalizations were recorded from 7 zebra finches housed in acoustic isolation chambers. On average birds were recorded from day 39 to 88 and live tutored from day 44 to day 67 post hatch. Audio was recorded continuously at a sampling rate of 32kHz and portions without vocalizations were discarded. Recorded audio was band-passed between 350Hz-8khz (using a digital filter).

### Segmentation into Syllables

The resulting band-pass filtered recordings were segmented based on RMS-amplitude of the signal. Let *w_t_* ∈ ℝ denote the band-passed raw audio signal. The RMS-amplitude of the signal at time *t* is defined as 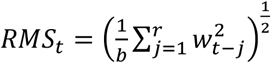. We chose the rms-buffer-size, *r*, to be 256 samples. Given a threshold 8 to separate signal from background noise, segment onsets were detected as upward threshold crossings of the RMS signal, i.e. an onset at time *t* requires that *RMS_t_*<*τ* and *RMS*_*t*+1_ ≥ *τ*,. Offsets are downward threshold crossings. Some segmentations were corrected in a semi-automatic process. We only corrected cases where both context (i.e. surrounding syllables) and spectral content clearly indicate a segmentation error.

### Spectrograms

Spectrograms of the detected segments were generated from the band passed raw audio signals. Spectrograms were computed using *B* = 512 samples (16ms) for each Fourier transform. A spectrogram column, *m_t_* ∈ ℝ^*B*^, is given by the discrete short time Fourier transform of a smoothed short section of the band-passed raw audio signal 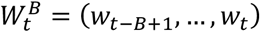. Smoothing is done by multiplying pointwise with a hamming smoothing window 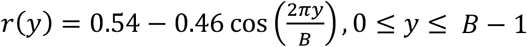. *m_t_* is computed at all timepoints *t* = *kS*) and we refer to *S* as the step-width of the spectrogram and *B* as the buffer size. For the analysis we used a step width of 64 samples.

We compute the absolute values of the Fourier coefficients of *m_t_* and keep only those coefficients corresponding to frequencies within the bandpass region. To provide increased amplitude invariance, we use log spectrograms and add an offset of 1:

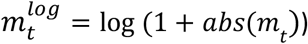

(all operations are pointwise). Several subsequent spectrogram columns 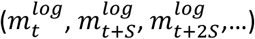 form one spectrogram snippet. For almost all analysis, snippets were 68ms long (2176 samples) and aligned to the onsets of vocalizations. For the large-scale t-SNE embeddings (e.g. Fig. 2A), 200ms (6400 samples) snippets were used. In those 200ms snippets, the subsequent syllable was masked if it overlapped with the snippet. For the example features in Fig. 1D-F, snippets of length 200ms (covering the entire syllable) were used.

We validated the stability of our recording conditions by studying RMS amplitude and embeddings of spectrogram snippets of background noise (i.e. when the bird is not singing).

The above described process yielded between 325,466 and 1,260,715 segments per bird of median length between 87 and 130ms. Totaled over all birds, this yields 6,227,626 data points, each with 3294 dimensions (121 frequency bins x 27 spectrogram columns) in the case of a 68ms window and 5734 dimensions in the case of a 200ms window.

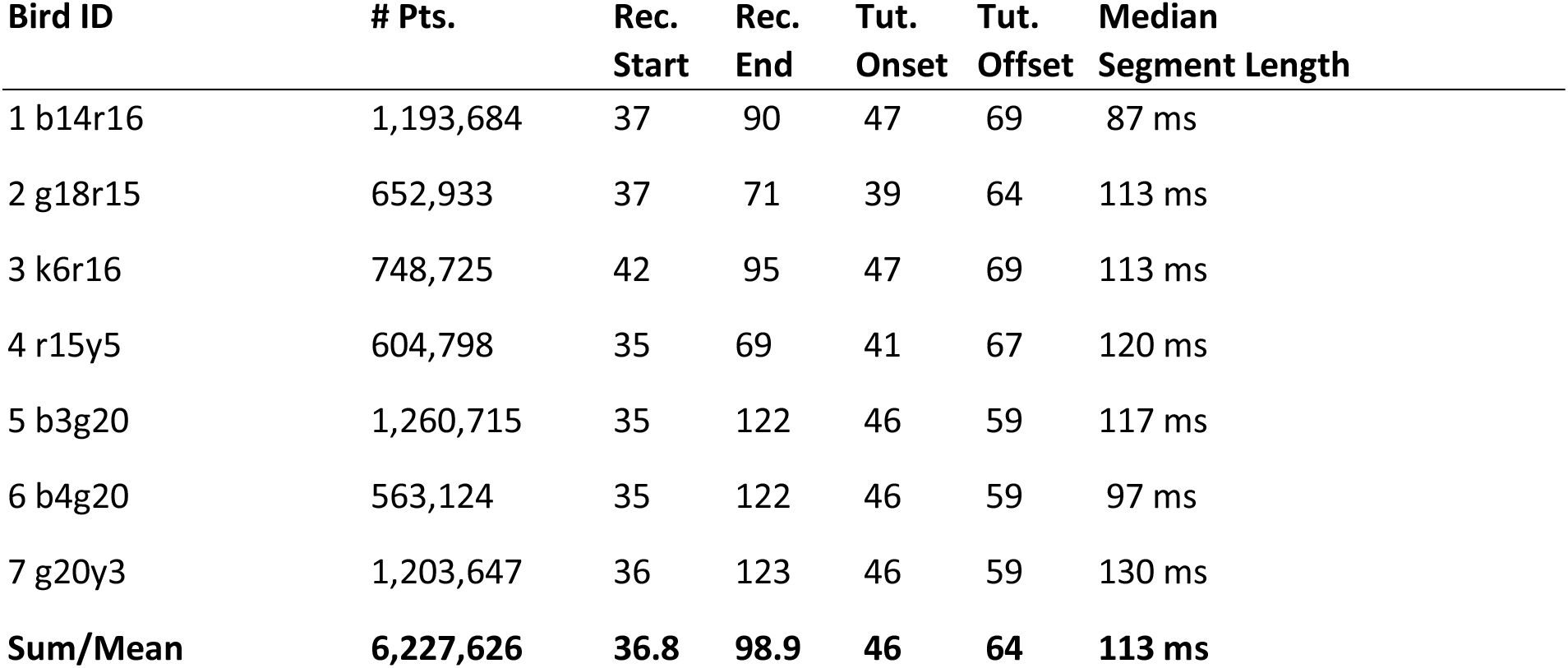

### Data used in individual figures

Figure 1: bird 6 (example)

Figure 2A-D: bird 6 (example)

Figure 2E, F: birds 1, 3, 5, 6 and 7

Figure 2G, H: bird 6

Figure 3A-D: bird 6

Figure 3E-K: birds 1, 3, 5, 6 and 7

Figure 4B-E: birds 1, 3, 5, 6 and 7

Ext. Data Fig. 1: bird 6

Ext. Data Fig. 3: birds 3, 5, 7

Ext. Data Fig. 4: birds 1, 3, 5, 6 and 7

Ext. Data Fig. 6: birds 1, 3, 5, 6, 7

Ext. Data Fig. 8: birds 1, 3, 5, 6, 7

Ext. Data Fig. 9: bird 1

Ext. Data Fig. 10A: birds 1-7

Ext. Data Fig. 10B, C: birds 5,6,7

This selection is based only on the length of continuous recordings available for each bird. For every analysis, we included all birds for which continuous recordings were available over the entire time period of interest.

### Large Scale Embedding and Sorting

To isolate song vocalizations, we first embedded all data up to and including day 80 post hatch using Barnes Hut t-SNE (described below).

Briefly, t-SNE ^1^ places data points in a 2 or 3 dimensional space, such that high dimensional local distances are approximately preserved. The algorithm computes a pairwise measure of data point similarity *p_ij_*. The measure is normalized and interpreted as a probability distribution. This distribution can be computed in the original high dimensional space as well as in the reduced low dimensional visualization space. By minimizing the KL-divergence between the distributions ascertained in these two spaces, a low dimensional visualization is generated that preserves structure determined by high dimensional local distances.

Large scale embeddings were computed on a high performance cluster using a custom version of Barnes-Hut-t-SNE ^2^. We used varying perplexities between 30 and 400. The embedding in Fig. 2 is computed using a perplexity of 200, with 5,000 iterations. The algorithm is iterative, and the solution depends on starting conditions. For embeddings with random starting conditions, we selected the best in terms of final KL-divergence after 5,000 iterations.

We used the embeddings to isolate calls, noise and song. Then we used nearest neighbor classification (10 nearest neighbors, majority vote, initial PCA to project data down to 150 dimensions) to classify data recorded after day 80. In a final step, we embed the combined song data for all days. We additionally separately embedded putative calls and noise post day 80 as identified by the nearest neighbor classifier to identify incorrect classifications.

Noise and isolated calls (i.e. calls not incorporated into song) were excluded from the analysis. All data recorded while several birds were in the isolation chamber (live tutoring) was excluded as well.

Based on the large scale embeddings and embeddings of all data generated on a single day, we separated the renditions into separate syllable types in a semiautomatic process. We use this clustering for visualization purposes (e.g. Extended Data Fig. 9), to perform a control based on local linear analysis (Extended Data Fig. 8) and to control for effects of syllable types in bout-based analyses (Fig. 3F, H, Extended Data Fig. 6).

### Bout Definition

Bouts are defined as sequences of consecutive vocalizations that are separated by pauses no longer than Δ*T* seconds. Δ*T* should reflect the “natural” temporal patterning of vocalizations. Values in the literature range from 160ms^3^, over 300ms^4^ to 2s^5^ or 30s^6^. A good Δ*T* should lead to small gaps between segments within a bout and large gaps between bouts. Our approach is to look at the problem as a clustering problem in ℝ^1^, where the data points are given by the segment midpoints in time. We assess the quality of the clustering by the Dunn index^7^ which measures how compact clusters are, compared to the distance between clusters:

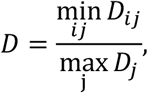

where *D_ij_* refers to the distance between centroids of clusters *j* and *j* (here the centroid is the average of all syllable mid points in the bout) and *D_j_* refers to the maximal distance between points in cluster *j* (here the separation between first and last syllable in a bout). A large Dunn index indicates dense well separated clusters. The Dunn index indicates that on average the best clustering is obtained with Δ*T*~2,500*ms* (not shown).

### Computing Nearest Neighbors

We use different methods to find nearest neighbors depending on dimensionality and data amount. For low dimensional data up to 10 dimensions, we use KD-Trees^8^. If the dimensionality of the data is higher, we use vantage point trees^9^ or exhaustive search. All nearest neighbor computations are exact. For high dimensional data, nearest neighbors are computed after projecting the data onto its first 100-300 principal components (250 principal components explained between 91% and 95% of the total variance) but results are reproducible for different numbers of principal components. Generally, we try to keep the number of nearest neighbors small to avoid contamination across clusters (compare Extended Data Fig. 1). However, we found our results to be robust over different numbers of nearest neighbors. Distance measure for identifying nearest neighbors is the standard Euclidean distance.

### Notation for Nearest Neighbor Statistics

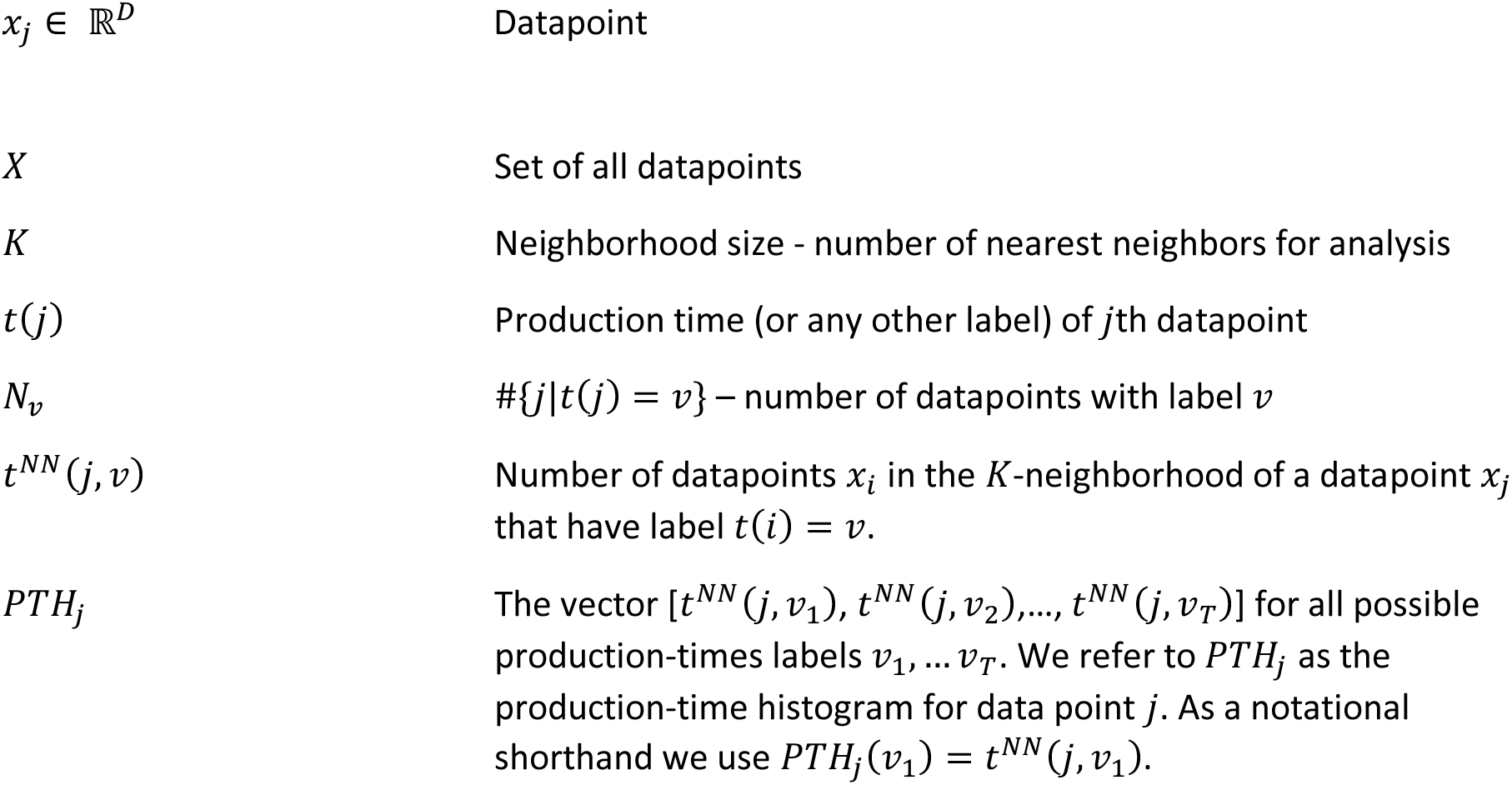

### Nearest Neighbor Histograms

Our approach for characterizing the structure of a dataset (for any sort of data and labels) is based on quantifying the composition of the neighborhood of each data point, whereby a neighborhood is defined as the set of *K* nearest-neighbors of a given data point. Specifically, we ask which labels are “locally close” to which other labels by computing statistics over the nearest neighbors of data points.

Consider a graph with data points as nodes and directed edges given by nearest neighbor connections (Extended Data Fig. 2B). Nodes *j* in this graph have labels given by *t*(*j*). The labels of a node’s neighbors can be counted. If labels are production times, we refer to those counts for a given datapoint as its *production time histogram*

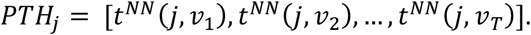

In the case of Fig. 2C, data points *x_j_* are spectrogram snippets and *t*(*j*) is the production time specified as day post hatch. For a datapoint *x_j_*, the value of the production time histogram *t^NN^*(*j,v*) for production time *v* is the number of neighbors of *x_j_* that have production time *t*(*i*) = *v*.

The production-time histogram is a special case of what we call a *nearest-neighbor histogram*, which can be computed in an analogous fashion for any arbitrary label *t*(*i*). For example, in Fig. 4 each data point is characterized by a label *t*(*i*) ∈ {1,…,50} indicating the bin to which the point belongs. Each of the 50 bins in turn is specified by a unique combination of stratum (1-5, regressions to anticipations), production period (1-5, morning to evening), and day (k or k+1). This binning is nested. First, we bin into days, then into production periods, and finally we bin into strata within each production period.

### Repertoire Dating

For a datapoint *x_j_* with index *j*. The *pseudo production day* is defined as the median of the set of all production times of the K nearest neighbors of datapoint *x_j_*.

For a set of *D* datapoints *Y* = {*x*_*i*_1__, *x*_*i*_2__,…, *x_i_D__*} ⊂ *X* with indices *IND_Y_* = {*i*_1_,…, *i_D_*} We define the *pooled neighborhood production time histogram* as the (pointwise) sum of all production time histograms of the datapoints in the set:

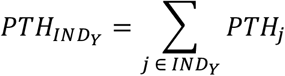

As before we refer to the summed count for label *v* as *PTH*_IND_Y__(*v*). The yth percentile (i.e. *P* = 50 for the median) of this pooled production time histogram is then defined through.

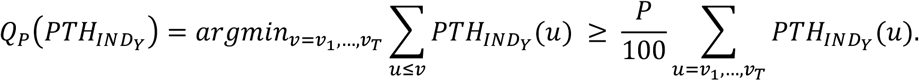

### Mixing Matrices

To summarize the neighborhood relations among all pairs of labels, nearest neighbor histograms can be added for all data points that carry a certain label. The resulting added counts summarize which labels are locally close to which other labels. A mixing matrix displays this information for all labels.

For a pair of labels *u* and *v* we define

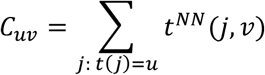

Since the number of data points can vary across production periods, we correct the raw counts *C_uv_* by dividing by the counts that would be expected if all neighborhoods were mixed. We define a null hypothesis for the mixing counts through:

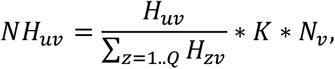

where *H_uv_* = *N_u_N_v_* and labels are assumed to come from the set {1..*Q*}, *K* is the neighborhood size and *N_v_* and *N_u_* denote the number of datapoints with label *v* or *u* (or *z*) respectively. We define the mixing matrix as the base-2-logarithm of the ratio of expected and observed counts:

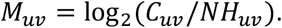

Each entry *M_vu_* of the mixing matrix assesses the similarity between data points with label *t*(*j*) = *v* and data points with labels *t*(*j*) = *v*. In particular, for the mixing matrix in Fig. 2D depicts on which days the bird produced similar syllables. Stationary behavior then results in blocks around the diagonal and non-stationarity is characterized by a concentration of large values onto the diagonal (Fig. 2). Note that all shown mixing matrices are computed in high-dimensional spaces based on spectrograms and Euclidean distances. Low-dimensional t-SNE embeddings (as in Fig. 2A) are only used for visualization purposes.

One notable property of mixing matrices is that they can be easily averaged across data sets (e.g. across birds, Extended Data Fig. 2E; or across pairs of days and birds; Extended Data Fig. 10). We average mixing matrices pointwise, irrespective of the number of data points entering the computation of each individual mixing matrix.

### Multi-Dimensional Scaling

To obtain a visual representation of the information represented in a mixing matrix, we use multidimensional scaling^10^. Each possible label *u* ∈ 1..*Q*, constitutes one data point to be placed in a low dimensional space and the mixing matrix defines the similarity structure among data points. Multidimensional scaling places these data points in a low dimensional space such that bins with high overlap (high value in the mixing matrix) are close-by. We perform non-classical multi-dimensional scaling, which matches only the order of dissimilarities but not their exact values.

In a first step we generate a dissimilarity matrix *D* from the mixing matrix *M* with elements *M_uv_* by setting

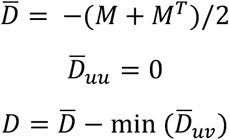

Second, we perform non-classical multi-dimensional scaling using the elements of *D* as input dissimilarities. Non-classical multidimensional scaling optimizes a low-dimensional representation that preserves the order of dissimilarities specified by *D*. Unlike t-SNE, multi-dimensional scaling does not focus on local distances. The algorithm is iterative, and solutions depend on starting conditions.

Non-classical multi-dimensional scaling computes a mapping between high and low dimensional dissimilarities that can be used to ascertain how well dissimilarity matrices are captured by the lowdimensional arrangement of data points (i.e. by the behavioral trajectory). The quality of the fit is assessed by a measure called ‘stress’ based on a residual sum of squares. As a rule of thumb, a stress value below 0.025 indicates an excellent fit and value below 0.05 a good fit; values around or larger than 0.2 indicate a poor fit^10^.

For visualization, the low-dimensional arrangement of points obtained with MDS is rotated such that axes are aligned with meaningful directions of change. In Fig. 2F and Extended Data Fig. 4, the axes correspond to the principle components of the inferred 10-dimensional behavioral trajectory. The axes of slow change in Fig. 2F correspond to the first two principle components. In Fig. 4C&D, we show projections along two axes of slow change, an axis of within-day change, and an axis of across-day change. To compute the axes of slow change, we averaged points over production periods and days, and computed the first two principle components of the resulting averages. To compute the axis of within-day change, we averaged points across strata and days, computed the direction spanning the first and last production time, and orthogonalized it with respect to the axes of slow change. To compute the axis of across-day change, we averaged points across strata and production periods, computed the direction spanning days k and k+1, and orthogonalized it with respect to the axes of slow change and within-day change. In Fig. 4E, the local direction of slow change is defined as the direction spanning two appropriate strata (see insets), after averaging points over production periods and days; the direction of within-day change is defined as in Fig. 4C, but orthogonalized with respect to the local direction of slow change.

### Local Linear Analysis

The linear example feature depicted in Fig. 1D-F, were computed as described below but without orthogonalization and on snippets of length 200ms.

The results in Extended Data Fig. 8, are obtained from a linear analysis performed separately for each syllable type. Data from the period 60-69 dph were considered as 68ms long, onset-aligned spectrograms. The analysis is performed for each window of 4 consecutive days contained within 60-69 dph (i.e. 60-63, 61-64, 62-65,…).

Let *X_k_* denote the set of all renditions for day *k* and *X_k_*,…,*X*_*k*+3_ the 4 days in an analysis window starting at day *k*. First, we subtract the mean across elements in *X_k_* ∪ … ∪ *X*_*k*+3_ from each element in *X_k_* ∪ … ∪ *X*_*k*+3_. Second, we fit a linear function, *f*(*x*) = *Am*, where *A* denotes a linear kernel of the same dimension as a 60ms spectrogram snippet, to the regression problem

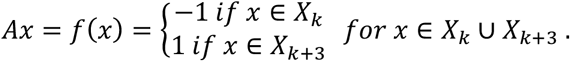

*f*(*m*) locally approximates the direction of slow change (Extended Data Fig. 8) as a 1d linear subspace spanned by *A*. Third, we fit a linear function *g*(*m*) = *Bm* to the regression problem

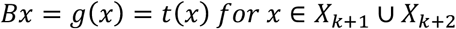

Where *t*(*x*) denotes the production period of *x* within the day (i.e. whether x was produced early or late). *g*(*m*) approximates within-day change by a one-dimensional subspace spanned by *B*.

Fourth, we fit a linear function *h*(*x*) = *Cx* to the regression problem

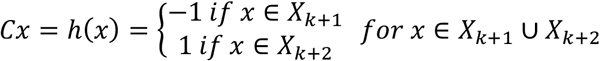

*h*(*x*) approximates across-day change by a one-dimensional subspace spanned by *c*. *f, g* and *h* were estimated using ridge regression with the regularization constant *λ* chosen through leave-one-out cross validation on the training set (Extended Data Fig. 8B). No data from days *X*_*k*+1_ and *X*_*k*+2_ was used to estimate *f*. No data from days *X_k_* and *X*_*k*+3_ was used to estimate *g*.

In a fifth step, we normalize <*A,B*> and C to norm 1 and orthogonalize *B* w.r.t. *A* through

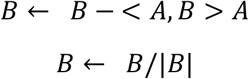

where < A,B > denotes the scalar product of *A* and *B* and |*B*| the norm of *B*. Finally, we orthogonalize *C* w.r.t. *A* and *B*.

Data from the inner days *X*_*k*+1_ and *X*_*k*+2_ is projected onto *A,B*, and *C*, and then averaged within each syllable of each bird. All syllables are then averaged again to obtain Extended Data Fig. 8D, E, F. Error bars depict 95% confidence intervals.

### Simulated Behavioral Development

We simulate individual behavioral renditions as points *r_i_* in a 100-dimensional state space (those correspond to *x_i_* above). Differences between renditions reflect (1) variability along a (time varying) piece-wise linear direction of slow change (DSC); (2) variability orthogonal to the DSC; and (3) high dimensional “shot noise”. The first two components of variability undergo distinct trends within a day and across days. Rendition *r_i_* ∈ ℝ^100^ is given by

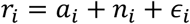

where *a_i_* ∈ ℝ^100^ determines the location along the DSC, *n_i_* ∈ ℝ^100^ a component orthogonal to the DSC, and *ϵ_i_* is normally distributed noise with identity covariance and a standard deviation of 0.001 along each dimension of *K*^100^. Each rendition *r_i_* is associated with a production time *t_i_*, given by *t_i_* = *d_i_* + *h_i_*, where *d_i_* = *floor*(*t_i_*) gives the day of production and 0 ≤ *h_i_* = *t_i_* – *d_i_* ≤ 1 gives the time within a day from early to late.

For each rendition, *r_i_*, the component *a_i_* is generated based on a “reference-time”

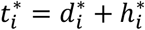

which corresponds to the time when renditions similar to rendition *r_i_* were typical in the behavioral repertoire. Note that in general 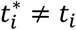 with 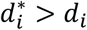 for visions and 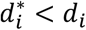 for regressions. By construction, the reference time closely matches the pseudo production-time for that rendition.

The reference time 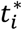 for a rendition *r_i_* produced at time *t_i_* is drawn from a time dependent probability distribution 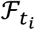:

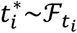

To account for differences in the time-course of visions and regressions, the distribution 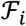 can be asymmetric. We refer to the part of 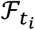 describing values larger than its median as the “positive” lobe, 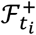, and to the part describing values smaller than its median as the “negative” lobe, 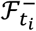. We define 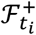 as the positive lobe of a (normalized) sum of two normal distributions with time dependent median *c*(*t_i_*) and standard deviations 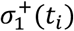 and 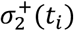. Likewise, 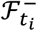 is defined as the negative lobe of a (normalized) sum of two normal distributions with the same median *c*(*t_i_*) and standard deviations 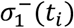 and 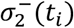. The, common, time-dependent median is given as

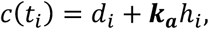

where ***k_a_*** determines the extent of within-day change along the DSC. Similarly, the time-dependent standard deviations are given by

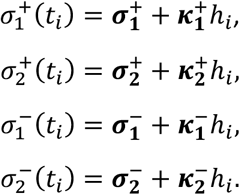

Where 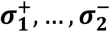 are scalar constants.

For each day we generate simulated behavior by first sampling 5000 production times *h_i_* uniformly from the unit interval [0,1]. To avoid edge effects, we simulated renditions for 240 subsequent days, and then analyzed only days 101 to 140 (Extended Data Fig. 5). Based the generated *t_i_,d_i_* and *h_i_* we sample reference times 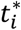 from 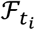 as described.

Having generated 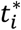 for rendition *r_i_*, we define the component along the DSC as:

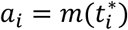

where *m*(*t*) ∈ ℝ^100^ is the DSC indexed by time *t*. The DSC is piece-wise linear, i.e. consists of a set of vertices and linear edges between subsequent vertices. The vertices occur at *h* = 0, meaning that vertex *m*(*d*) corresponds to song typical at the beginning of day *d* and the edge between *m*(*d*) and *m*(*d* + 1) describes the progression of song from morning to evening on day *d*. We define the local direction *v_d_* of the DSC as *v_d_* = *m*(*d* + 1) – *m*(*d*). We initialize *m*(1) and *v*_1_ randomly and then define the position of each subsequent vertex iteratively as:

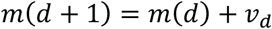

We impose |*m*(*d*)| = 1. The local direction of the manifold *v_d_* is chosen randomly under the constraints |*v_d_*|= 0.05 and *angle*(*v*_1_, *v*_1_) = 10°. For an arbitrary time *t* = *d*+*h* the DSC is defined as:

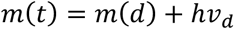

Notably, for all *d, m*(*d*) and *v_d_* are chosen from within a fixed, random, 90-d subspace of ℝ^100^, whereas the component *n_i_* orthogonal to the DSC is chosen within the 10-d subspace orthogonal to this 90-d subspace. Specifically, *n_i_* is drawn from a 10-d normal distribution 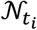:

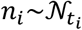

The distribution has isotropic standard deviation of 2 and a time depended mean *z_t_i__* given by:

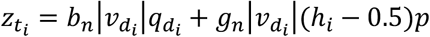

*q_d_* is a unitary vector specifying the direction of *across-day* change orthogonal to the DSC and is chosen randomly for each day *d* within the 10-d subspace; *p* is a random unitary vector specifying the direction of *within-day* change orthogonal to the DSC and is fixed across days; the parameters *b_n_* and *g_n_* determine the size of across-day and within-day components of change orthogonal to the DSC.

We simulated data based on two set of parameters that correspond to different relative alignments between the DSC and the direction of within-day and across-day change (Extended Data Fig. 5). The parameters were chosen by hand to qualitatively match the range of within-day and across-day variability observed in the data. The observed match between the outcome of repertoire dating (Extended Data Fig. 5C, D) and the ground truth (Extended Data Fig. 5A, B) is robust to changes in the simulation parameters.

#### Model 1 – weak consolidation

1. *b_n_* = 1, i.e. the component of across-day change orthogonal to the DSC is as large as the component along it;
2. *g_n_* = 0, i.e within-day change has no component orthogonal to the DSC;
3. *k_a_* = 5, i.e. the component of within-day change along the DSC corresponds to 5 times the amount of across-day change along the manifold, implying weak consolidation;
4. 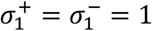 and 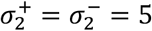
5. 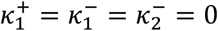 and 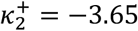, i.e. visionary renditions move less far along the DSC within a day compared to typical renditions and regressions.

#### Model 2 – strong consolidation

1. *b_n_* = 0, i.e. across-day change has no component orthogonal to the DSC;
2. *k_a_* = 1.25; i.e. the component of within-day change along the DSC is only slightly larger than the component of the across-day change along it, implying strong consolidation;
3. *g_n_* = 2; i.e. the largest component of within-day change is orthogonal to the DSC;
4. 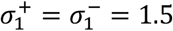 and 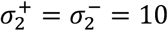
5. 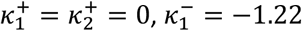, and 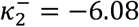, i.e. regressive renditions move farther along the DSC within a day compared to typical renditions and visions.

### Ascertaining Clustering Structure

In Extended Data Figure 1 we show daily t-SNE embeddings of 60 ms vocalization snippets as well as estimates of the number of clusters in the data based on k-means clustering. The emergence of clusters is apparent in the t-SNE visualizations. We reproduced this finding directly on the high dimensional data by performing k-means on the data of each day and varying cluster numbers. We compare different k-means clustering solutions using silhouette coefficients^11,12^. These cluster validity indices are computed as follows.

For each datapoint *x_i_* and each found cluster *C_k_* ⊂ *X* we define

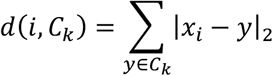

to be the average distance of *x_i_* to each element in *C_k_*. Let *x_i_* be in cluster *C_k_*. We define the average distance of *x_i_* to points in its own cluster *C_k_*:

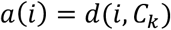

and the minimum average distance of *x_i_* to points from another cluster:

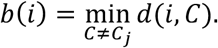

The silhouette coefficient is then defined as

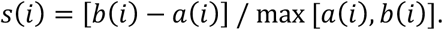

This definition requires at least 2 clusters and at least 2 datapoints per cluster. *s*(*x_i_*) ranges from −1 to 1. A silhouette coefficient of −1 indicates that a datapoint is not well ‘embedded’ with its own cluster whereas +1 indicates that it is. As a measure for the overall quality of a clustering we use the average of *s*(*i*) over all datapoints^12^.

## EXTENDED DATA FIGURES

**Extended Data Figure 1.**
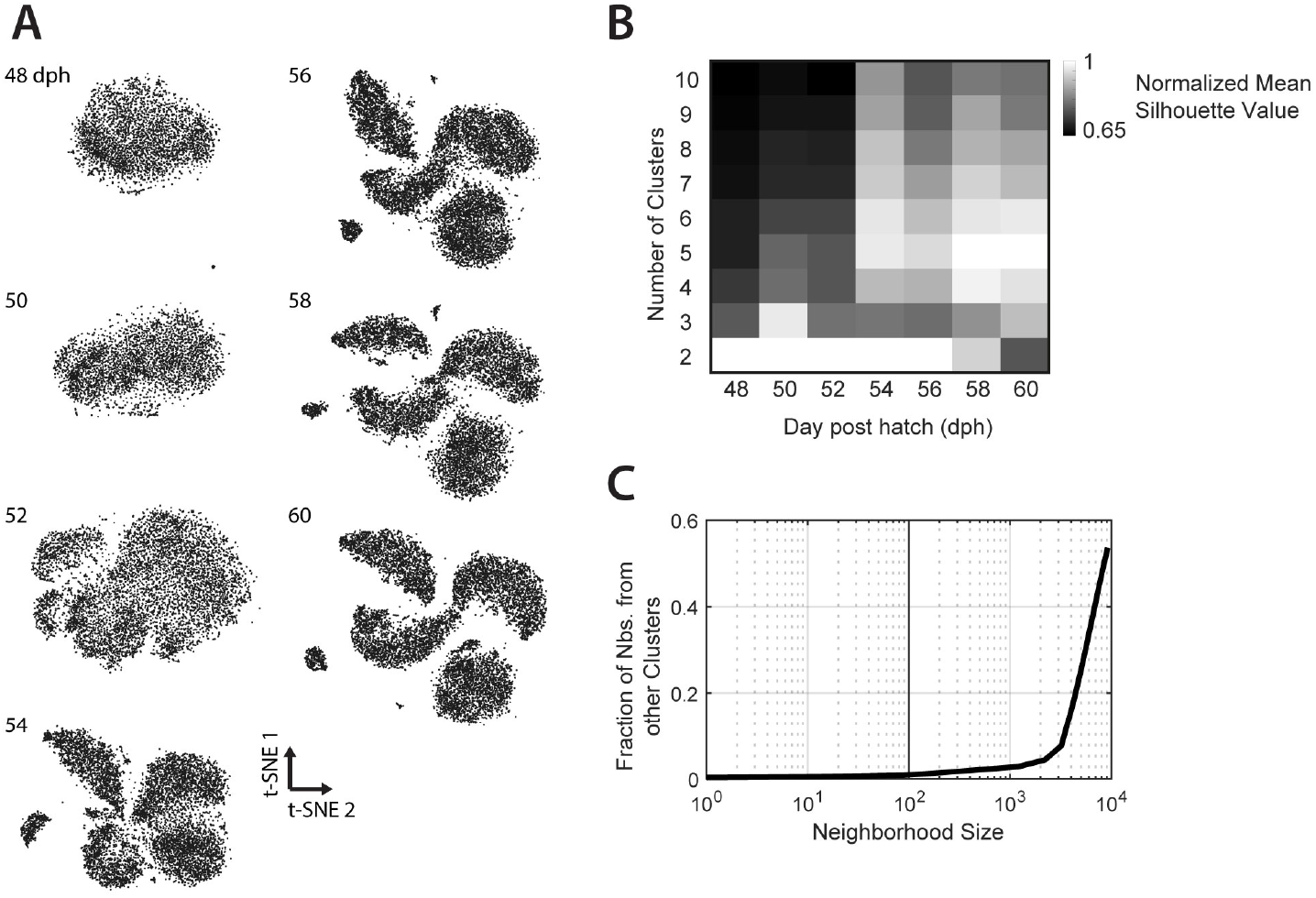
Clustering of juvenile and adult Zebra finch song. (**A**) t-SNE embeddings of vocalizations produced on the specified days (labels; dph) for the example bird depicted in Fig. 1 and 2A-D. Here vocalizations were represented as 60ms onset-aligned spectrogram snippets and a separate embedding was computed for each day. Note the gradual emergence of clusters, each corresponding to a distinct syllable type (e.g. syllables i, a, b, c in Fig. 2A). (**B**) Normalized mean silhouette values (see Methods) for 2-10 clusters computed on vocalizations from the 7 days in (A). A high value in a cell of this matrix indicates that there is evidence for the respective cluster count in the vocalizations from the corresponding day. Mean normalized silhouette coefficients were computed based on 20 repetitions of k-means clustering of 60ms onset-aligned spectrogram snippets from a single day (same as in A) projected onto the first 50 principal components. (**C**) Average fraction of neighbors from a different cluster, as a function of neighborhood size. Analogous data to (A) and (B) but for vocalizations from dph 90 (12,854 data points), when clusters are fully developed. For a wide range of neighborhood sizes, the neighbors of a data point mostly belong to the same cluster or syllable type. For a neighborhood size of 100, the average fraction of out-of-cluster neighbors from the same day is 0.0089. This implies that, for an appropriately chosen neighborhood size, nearest-neighbor methods respect clustering structure in the data by construction. Nearest-neighbor statistics thus sidestep having to explicitly identify clusters in the data. Note that in most analyses we computed nearest-neighbors on data from all days, meaning that clustering structure is respected even for neighborhood sizes somewhat larger than those implied in (C).

**Extended Data Figure 2.**
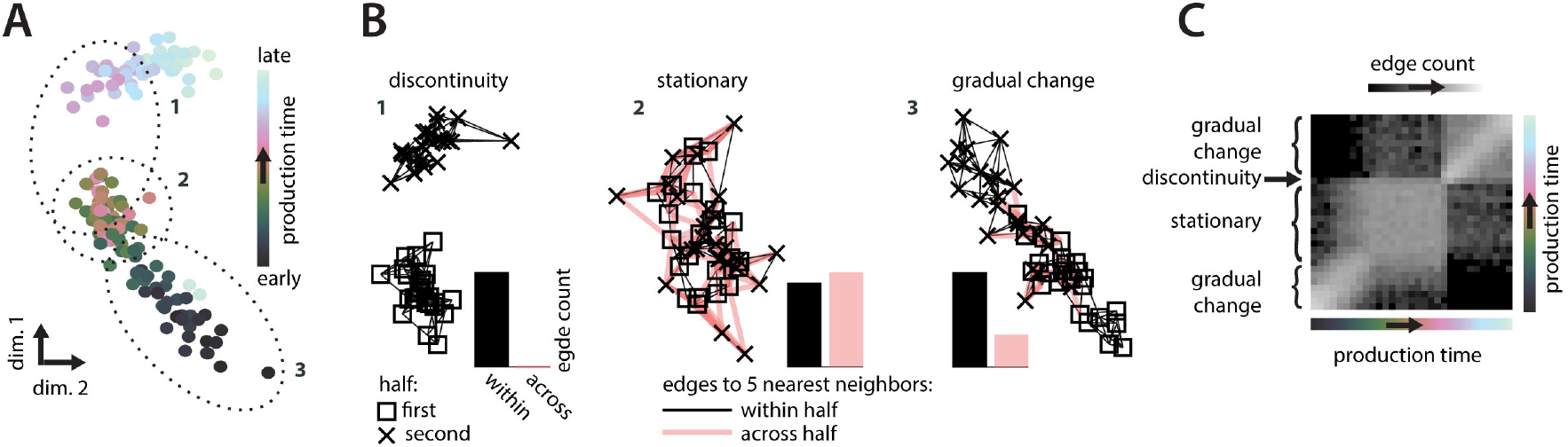
Characterizing behavioral change using neighborhood graphs. Illustration of a fictitious behavior undergoing distinct phases of gradual change, no change, and abrupt change, and the identification of these phases based on nearest-neighbor graphs. (**A**) A low-dimensional representation of the behavior. Each point corresponds to a behavioral rendition (e.g. a syllable rendition in the bird data) and is colored based on its production time. Similar renditions (e.g. that have similar spectrograms) appear near each other in this representation. The dotted ellipses mark three subsets of points corresponding to (1) a phase of abrupt change; (2) a stationary phase; and (3) a phase of gradual change. (**B**) Nearest neighbor graphs for the three subsets of points in (A). Points are replotted from (A) with different symbols, indicating whether their production times fall within the first (squares) or second half (crosses) of the corresponding subset. Edges connect each point to its 5 nearest neighbors. Edge color marks neighboring pairs of points falling into the same (black) or different (red) halves. Relative counts of within- and across-half edges differ based on the nature of the underlying behavioral change (histograms of edge counts). If an abrupt change in behavior occurs between the first and second half, nearest neighbors of points in one half will all be points from the same half, and none from the other half (discontinuity). When behavior is stationary, the neighborhoods are maximally *mixed*, i.e. every point has about an equal number of neighbors from the two halves (stationary). Phases of gradual change result in intermediate levels of mixing (gradual change). (**C**) Nearest-neighbor mixing-matrix for the simulated data in (A). Each location in the matrix corresponds to a pair of production times. Strong mixing (white) indicates a large number of nearest-neighbor edges across the two corresponding production times (as in B, stationary) and thus similar behavior at the two times. Weak mixing (black) indicates a small number of such edges (as in B, discontinuity), and thus dissimilar behavior. Note that such statistics on the composition of local neighborhoods can be computed for any kind of behavior and are invariant with respect to transformations of the data that preserve nearest neighbors, like scaling, translation, or rotation. These properties make nearest-neighbor approaches very general.

**Extended Data Figure 3.**
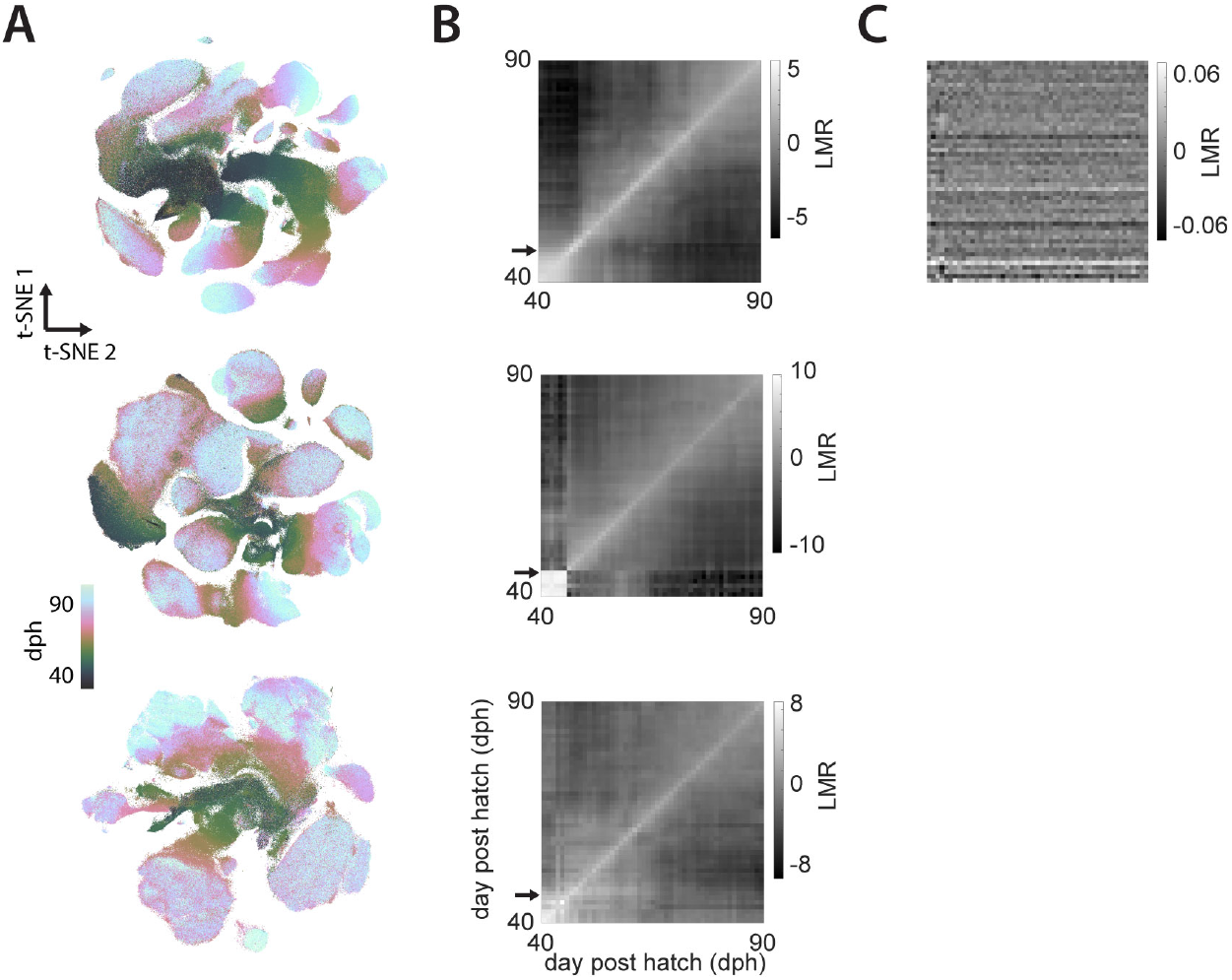
t-SNE embeddings and mixing matrices for 3 additional birds. (**A**) t-SNE embeddings for the continuous recordings from 3 example birds (rows). Analogous to Fig. 2A. (**B**) Lifetime mixing-matrices for the data in (A). Analogous to Fig. 2D. Black arrows indicate tutoring onset. The bird in the middle row produced only very few vocalizations (mostly calls) before tutoring onset. The mixing-matrices consistently show a period of gradual change starting after tutor onset and lasting several weeks. Gradual change typically slows down (resulting in larger mixing values far from the diagonal, e.g. bottom bird) but is still ongoing at the end of the developmental period considered here (dph 90). Gray values correspond to the base-2 logarithm of the mixing ratio, i.e. counts in the pooled neighborhood production-time histograms (Fig. 2C) normalized by a null-hypothesis obtained from a random distribution of production times (see Methods). For example, an LMR value of 5 implies that renditions from the corresponding pair of production-times are 2^5^ = 32 times more mixed at the level of local neighborhoods than expected by chance (i.e. a random distribution of production times across renditions). (**C**) Like (B), top, but after shuffling production times among all data points. Effects under this Null-hypothesis are small (the maximal observed mixing ratio is 2^0.06^ ~ 1.042). Similar, small effects under the Null-hypothesis are obtained for the other mixing matrices discussed throughout the text.

**Extended Data Figure 4.**
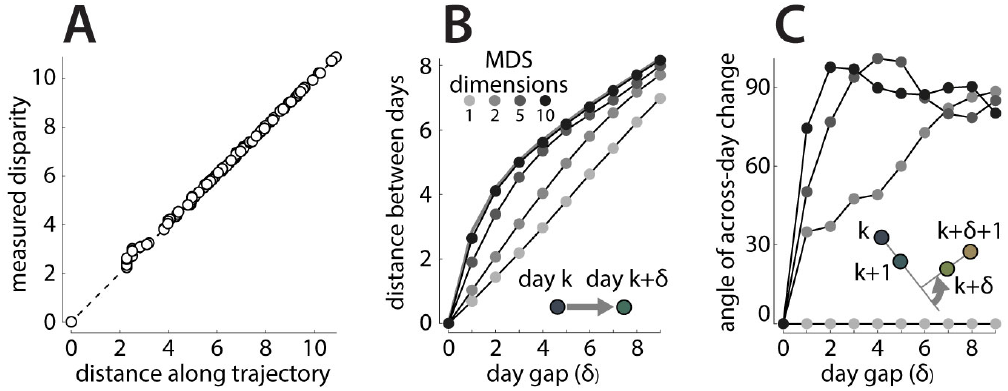
Structure of change along the inferred behavioral trajectory. Properties of the behavioral trajectory inferred from the mixing-matrix in Fig. 2E. (**A**) Pairwise distances between points along the inferred behavioral trajectory (horizontal axis) compared to the measured disparities (vertical axis). Disparities are obtained by rescaling and inverting the similarities in Fig. 2E (see Methods). The points on the trajectory are inferred with 10-dimensional non-metric multidimensional scaling (MDS) on the measured disparities. Crucially, the pairwise distances between inferred points faithfully represent the corresponding, measured disparities (all points lie close to the diagonal; MDS stress = 0.0002). (**B-C**) Structure of lowdimensional projections of the behavioral trajectory. We applied principle component analysis to the 10-d arrangement of points inferred with MDS and retained an increasing number of dimensions (number of dimensions indicated by grayscale) to compute low-dimensional projections of the full 10-d behavioral trajectory. For example, the projection onto the first two principle components is shown in Fig. 2F (MDS dimensions 2 in (B) and (C)). The first two principle components explain 75% of the variance in the full 10-d trajectory. (**B**) Measured disparity (thick gray curve) and distances along the trajectories (points and thin curves) as a function of the day-gap d between points. For any choice of projection dimensionality and d, we computed the Euclidian distances between any two points separated by d and averaged across pairs of points. The measured disparities increase rapidly between subsequent and nearby days, but only slowly between far apart days (thick gray curve). Low dimensional projections of the trajectory (e.g. MDS dimensions 2) underestimate the initial increase in disparities. (**C**) Angle between the reconstructed direction of across-day change for inferred behavioral trajectories, as a function of the day-gap between points. Same conventions and legend as in (B). For the 1-d and 2-d trajectories, the direction of across-day change varies little or not at all from day to day (see inset, arrow indicates angle of across-day change). On the other hand, the direction of across-day change along the full, 10-d behavioral trajectory is almost orthogonal for subsequent days. Both (B) and (C) imply that the full behavioral trajectory is more “rugged” than suggested by the 2d-projection in Fig. 2F. This structure is consistent with the finding that across-day change includes a large component that is orthogonal to the directions of slow change and of within-day change (Fig. 4C, bottom).

**Extended Data Figure 5.**
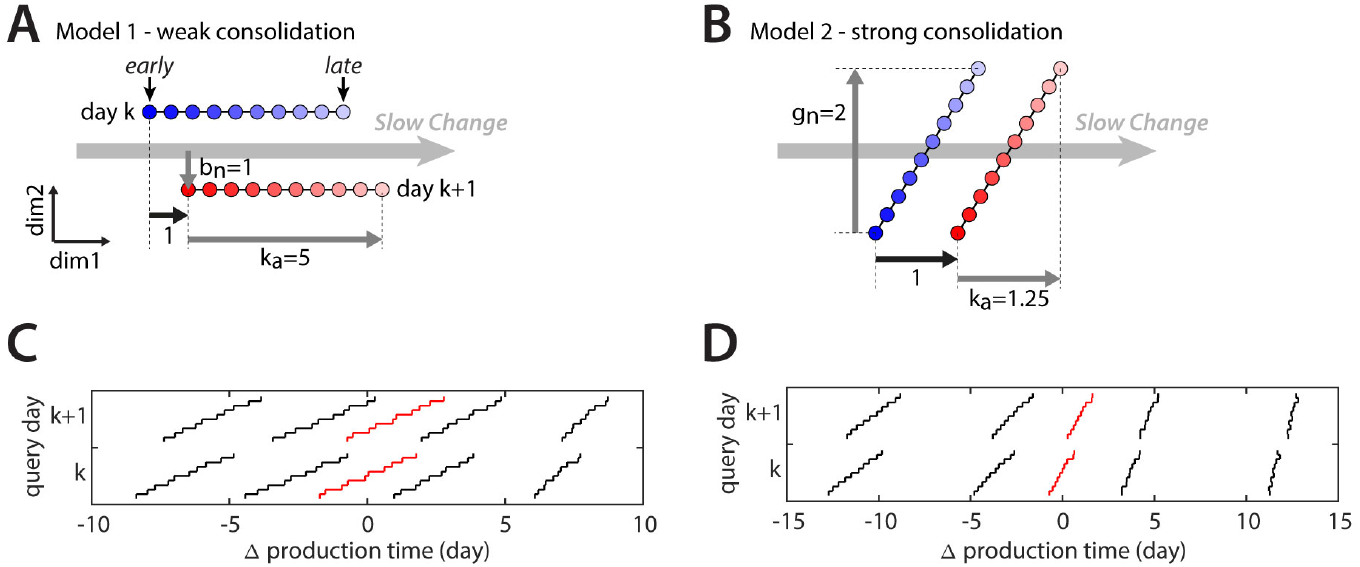
Models of behavioral change.

We simulated individual behavioral renditions as points in a high-dimensional space, drawn from a time-dependent probability distribution changing both within and across days (see Methods) and verified that repertoire dating (Fig. 3) can successfully recover the underlying structure of the models. The main parameters determining the relative alignment of the direction of slow change (*DSC*) with the directions of within-day and across-day change are ***k_a_***, the amount of within-day change along the DSC; ***g_n_***, the amount of within-day change orthogonal to the DSC; and *b_n_*, the amount of across-day change orthogonal to the DSC. These parameters are expressed relative to the amount of across-day change along the DSC (thick black arrow in (A) and (B)). The two models shown (see Methods) imply different amounts of overnight consolidation of within-day changes along the DSC. (**A**) Schematic illustration of model 1. Within-day change is aligned with the DSC (*g_n_* = 0) and large, i.e. corresponding to the overall distance traveled along the DSC over 5 days (*k_a_* = 5). The component of across-day change orthogonal to the DSC is as large as the component of across-day change along it. In this scenario, overnight consolidation of within-day changes along the DSC is weak (20% of change is consolidated) for typical renditions. (**B**) Schematic illustration of model 2. Within-day change has a large component orthogonal to the DSC, whereas across-day change is aligned with the DSC. In this scenario, overnight consolidation of within-day changes along the DSC is strong (80% of the change is consolidated) for typical renditions. (**C**) Repertoire dating results for model 1, analogous to Fig. 3G. Lines from left to right denote 5^th^, 25^th^, 50^th^ (red line), 75^th^ and 95^th^ percentile of the pooled neighborhood production times. Dating of the typical renditions (red) closely reproduces dynamics of change along the DSC implied by (A)—within-day change along the DSC is large (red line extends over about 5 days) and consolidation is weak (starting point on day k+1 relative to day k moves by about 20% of overall within-day range). Differences between visions (95^th^ percentile) and regressions (5^th^ percentile) correctly reflect the underlying model parameters (see Methods). (**D**) Repertoire dating results for model 2, analogous to (C). Dating of the typical renditions (red), visions and regressions closely reproduces dynamics of change along the DSC implied by (B).

**Extended Data Figure 6.**
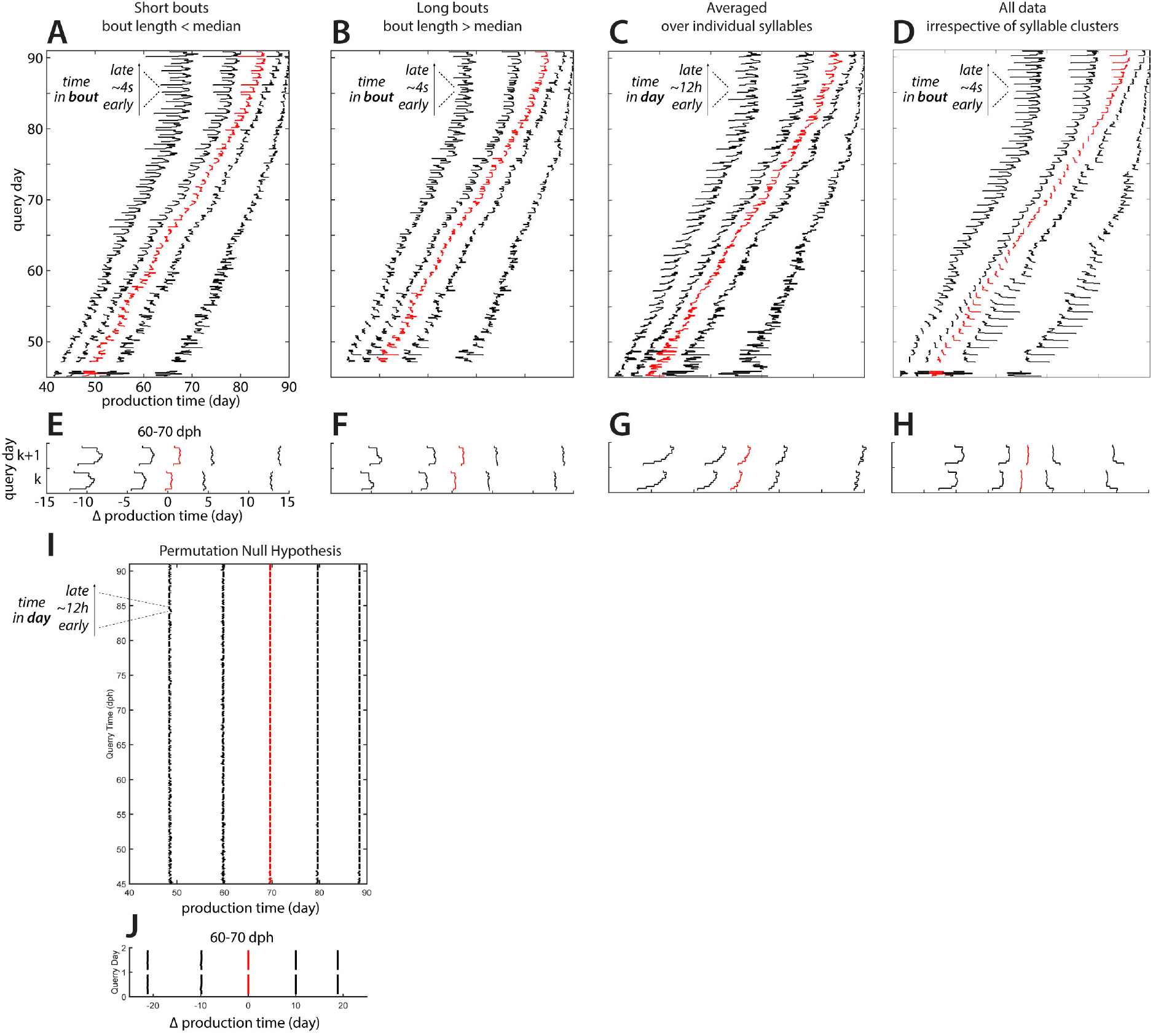
Repertoire dating control analyses. (**A**) Analogous to Fig. 3F, but computed only from renditions falling into short bouts (bout length < median). (**B**) Analogous to Fig. 3F, but computed only from renditions falling into long bouts (bout length > median). (**C**) Analogous to Fig. 3E, but computed for individual syllables, and then averaged across syllables and animals. Syllable clustering is described in the Methods. (**D**) Analogous to Fig. 3F, but computed over the entire data set without prior clustering into syllables. (**E**) Analogous to Fig. 3H, but based on (A). (**F**) Analogous to Fig. 3H, but based on (B). The changes in the behavioral repertoire observed within a bout are qualitatively similar for short and long bouts (compare E and F). In particular, the song becomes more regressive shortly before the end of a bout (5^th^ percentile, left-most curves). The analogous effect in Fig. 4H thus occurs at the end of a bout and not at a particular time after the beginning of a bout. (**G**) Analogous to Fig. 3G, but based on (C). The changes in behavioral repertoire are qualitatively similar to those in Fig. 3G, which are computed without prior clustering of syllables. This similarity implies that the dynamics along the direction of slow change in Fig. 3 cannot be explained by changes in the frequency of syllables sung during the day. (**H**) Analogous to Fig. 3H, but based on (D). The changes in behavioral repertoire are mostly similar to those in Fig. 3H, which are computed on individual syllables and then averaged across syllables. One difference with Fig. 3H are pronounced within-bout changes for anticipations early during development. These changes occur when neighborhoods are computed on the un-clustered data, and thus appear to be explained by changes in the frequency of syllables (e.g. introductory notes) sung throughout a bout. (**I**) Analogous to Fig. 3E, but computed after shuffling production times among all data points. Within-day changes of the percentile curves are small under this null hypothesis. (J) Analogous to Fig. 3G, but computed from (**I**). The maximal span of within-day fluctuations is 0.2 days, compared to 3.71 for the unshuffled data in Fig. 3G. The total repertoire spread (5^th^ – 95^th^ percentile) is around ~40 days compared to ~23 days for unshuffled data.

**Extended Data Figure 7.**
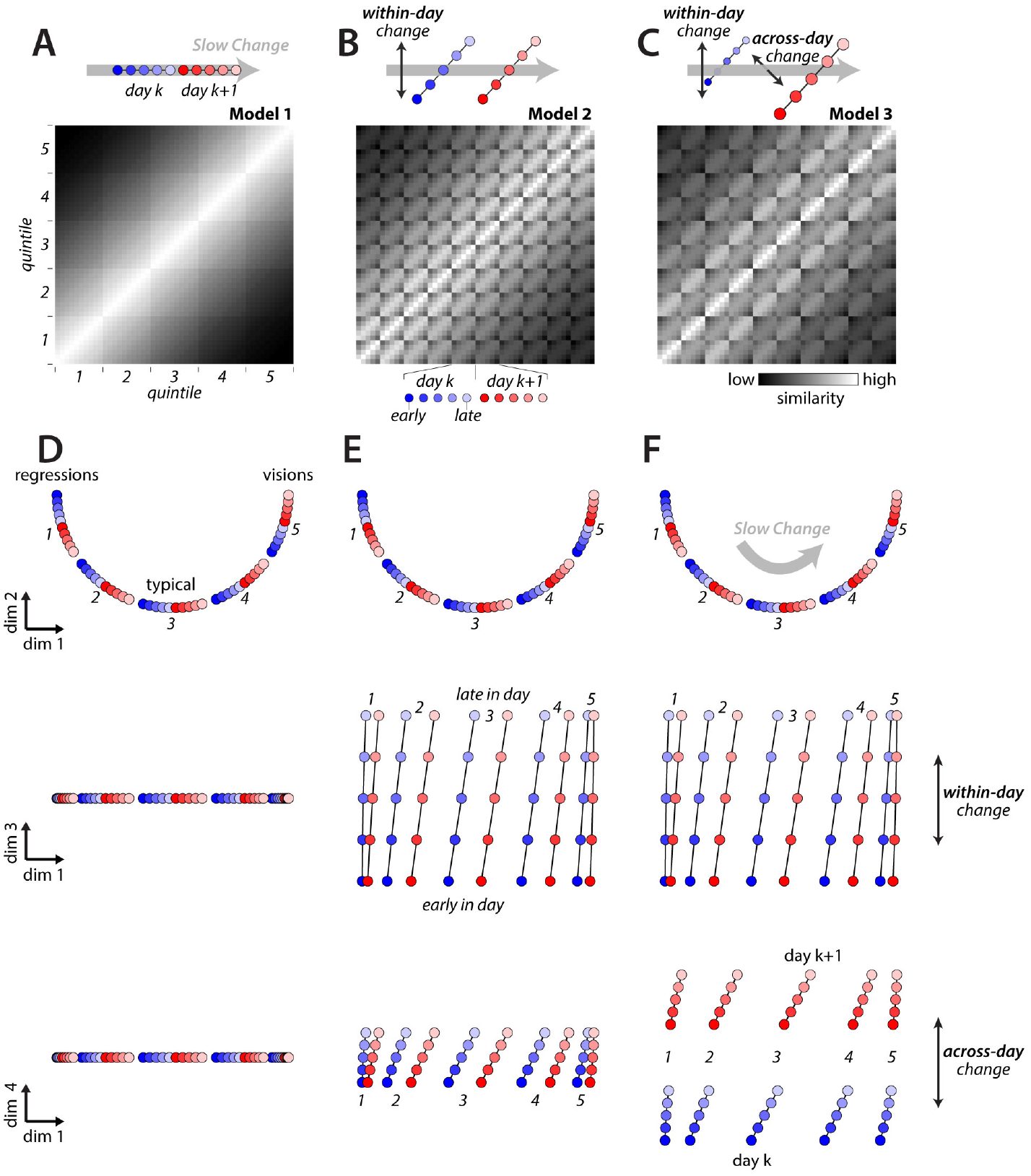
Alignment between the direction of slow change and change on short time-scales. We generated 3 sets of stratified behavioral trajectories (see also Fig. 4) that differ with respect to the alignment of within-day and across-day change with direction of slow change (*DSC*). We build each set of trajectories by arranging 50 points (5 strata per day, 5 production-time periods per day, on 2 consecutive days; same conventions as Fig. 4C) within a 4-dimensional space. We then generate simulated stratified mixing matrices (A-C, replotted from Fig. 4A) by computing pairwise distances between all points, and transforming distances into similarities. We visualize the behavioral trajectories (D-F) with the same 2-d projections as in Fig. 4C, with the same scale along all dimensions. In all models, overnight consolidation along the DSC is perfect for all strata. (**A**) Model 1: within-day change and across-day change occur only along the DSC. For each stratum (i.e. each of the five 10-by-10 squares along the diagonal) similarity decreases smoothly with time, reflecting the gradual progression of the trajectory along the DSC within and across days. (**B**) Model 2: within-day change has a large component that is not aligned with the DSC. For each stratum, song early on day k+1 is more similar to song early on day k, rather than to song late on day k as in model 1 (A). (**C**) Model 3: both within-day and across-day change have large components that are not aligned with the DSC. The misaligned component of across-day change reduces the similarity between day k and day k+1 compared to model 2, resulting in smaller values in the 5-by-5 squares comparing points from day k and day k+1. (**D**) Behavioral trajectories for model 1: the 2-d projection containing the DSC (top) explains all the variance in the trajectories. (**E**) Behavioral trajectories for model 2: similar to (D), but points from different periods during the day are displaced also along an orthogonal direction of within-day change (middle). (**F**) Behavioral trajectories for model 3: similar to (E), but points from adjacent days are displaced also along an orthogonal direction of across-day change (bottom). Note that models 1 and 2 in (A) and (B) are distinct from models 1 and 2 in Extended Data Fig. 5 (see Methods).

**Extended Data Figure 8.**
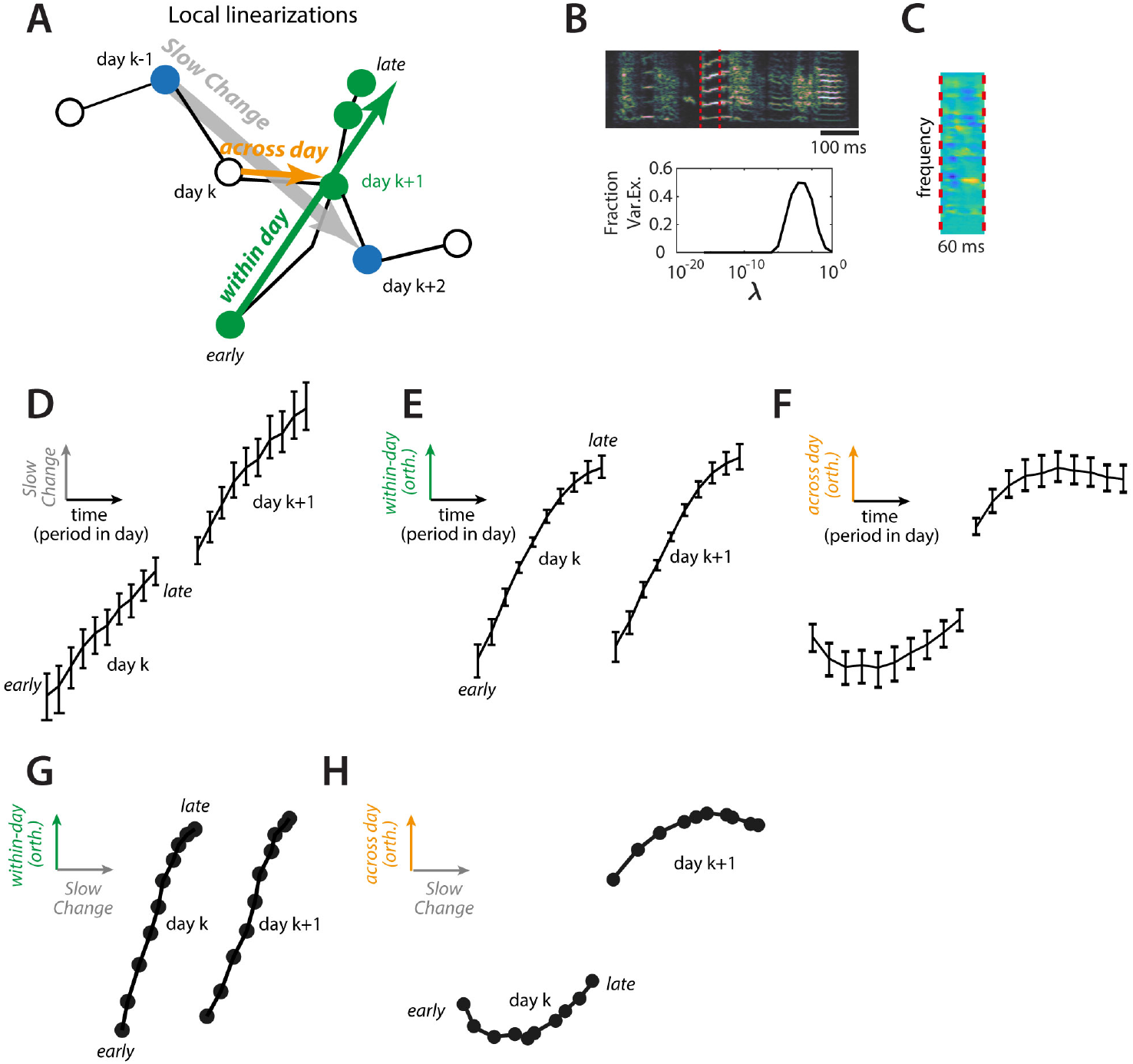
Local linear analysis. We validated the structure of change inferred with nearest-neighbor statistics (Fig. 4) with a more conventional approach based on linear regression in the high-dimensional spectrogram space (see Methods). Unlike for the case of nearest-neighbor based statics, here each rendition must first be assigned to a cluster (i.e. a syllable, compare Fig. 2A) and each cluster is analyzed separately. (**A**) Illustration of the linearization scheme. We infer the (local) direction of slow change (DSC) on day k (gray arrow) as the vector of linear-regression coefficients relating production day to variability in the renditions from days k-1 and k+2. Likewise, to infer the direction of within-day change (green arrow) we find linear-regression coefficients relating the period within a day to variability of renditions from days k and k+1 and then orthogonalize the coefficients with respect to the DSC. Both sets of coefficients, and the corresponding directions in spectrogram space, typically vary across days and syllables. The progression of song along the DSC and along the (orthogonalized) direction of within-day change is obtained by projecting renditions on day k and k+1 onto the corresponding (normalized) directions (see Methods). (**B**) Example rendition (top; encapsulated by red lines) and the dependency of crossvalidated regression quality (fraction of variance explained) on the regularization constant for the estimation of the DSC. One regularization constant was chosen for each syllable and direction based on maximizing the leave-one-out cross validation error on the training set. (**C**) Regression coefficients corresponding to the DSC for the example syllable in (B), estimated for day k = 65. Warm and cold colors mark spectrogram bins for which power increases or decreases, respectively, between days k-1 and k+2 (i.e. days 64 and 67). (**D**) Progression of song along the DSC as a function of time. Renditions from each day are binned into 10 consecutive periods based on production time within the day (analogous to the 10 periods in Fig. 3E). Projections onto the DSC for each period of days k and k+1 are then averaged across all 4-day windows during dph 60-69 and averaged again over all syllables and birds (same 5 birds as in Figs. 3, 4). The resulting averages include contributions from regressions, typical renditions, and visions. The position along the DSC for the morning of day k+1 is close to that for the evening of day k, indicating overall strong consolidation (Fig. 1D). Note that for simplicity of visualization, the time elapsed (horizontal axis) during the night between days k and k+1 is not shown. (**E**) Progression along the direction of within-day change as a function of time, analogous to (D). The position along the direction of within-change is reset overnight, implying that the underlying changes are not consolidated (Fig. 1E). By considering a feature that is a mixture of within-day change (E) and change along the DSC (D) a wide range of apparent consolidation scenarios could be “uncovered”. (**F**) Progression along the direction of across-day change as a function of time, analogous to (D). The position along the direction of across-day change jumps overnight. (Fig. 1F). **(G)** Progression along the DSC and the direction of within-day change, combining data from (D) and (E). This representation is analogous, and in qualitative agreement, with the behavioral trajectories in Fig. 4E (typical). (**H**) Progression along the DSC and the direction of across-day change, combining data from (D) and (F). This representation is analogous, and in qualitative agreement, with the behavioral trajectories in Fig. 4C (typical).

**Extended Data Figure 9.**
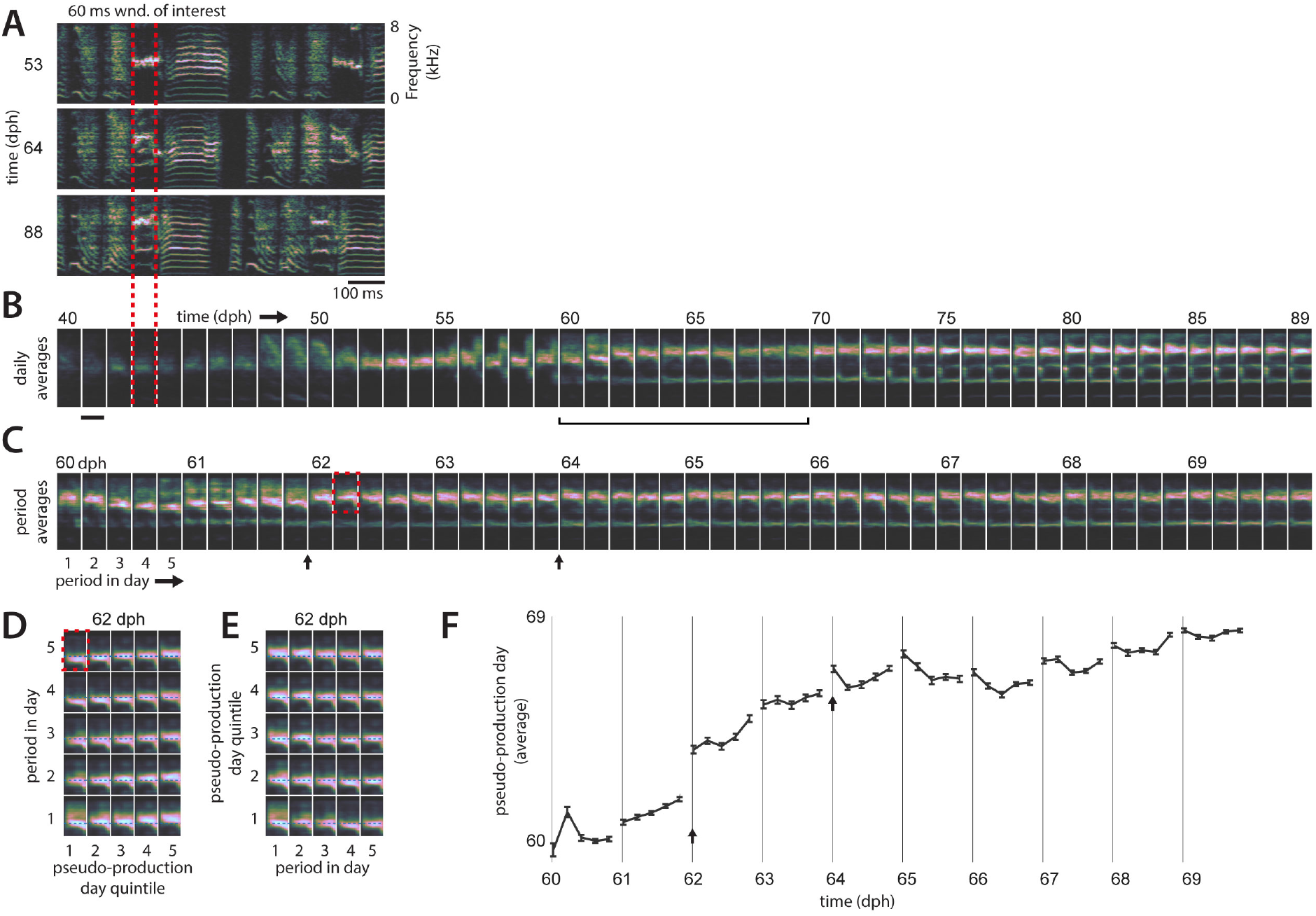
Behavioral variability and stratification in an example syllable. (**A**) Example songs of an example bird for 3 days during development. Vocalizations in the window of interest (red dotted lines, 60 ms) are analyzed in the following panels. (**B**) Developmental changes over the course of weeks. Vocalizations are binned by production day, and averaged. The most apparent changes occurring during learning are an increase in pitch and the later, successive appearance of additional spectral lines at low frequencies. (**C**) Within-day and across-day changes for the period 60-69 dph. Vocalizations are binned into 5 period spanning a day and averaged. On many days, the changes within a day do not appear to recapitulate the changes occurring across days (e.g. days 60 and 65, compare to subsequent days in (B). The averages also reveal occasional overnight “jumps” in the properties of the vocalizations (e.g. black arrows). (**D**) Comparison of within-day and changes on longer times-scales. Renditions within each period within a day were split into strata according to their pseudo-production day (quintiles in Fig. 3C) resulting in 25 averages, one for each combination of stratum and period within the day. Only the upper part of the spectrogram is shown (red rectangle in C). The progression along strata (x-axis) emphasizes the large extent of motor variability along the DSC existing within a single day (day 62). (**E**) Same averages as in D, but with x and y axes swapped. In particular for regressive renditions (quintile 1), change within day 62 (x-axis) does not recapitulate developmental changes occurring over months (x-axis in D). (**F**) Repertoire dating based on the pseudo-production day (as in Fig. 3C). Each point corresponds to a production-time period and is the mean of all pseudo-production days of renditions in that period. Errorbars show bootstrapped 95% confidence intervals. The change in pseudo-production day, which is computed without using any explicit spectrogram features, captures the movement along the DSC. Some of the effects seen in the raw spectrograms (B-E), including jumps and a pattern of within-day change that does not recapitulate the slow developmental changes (e.g. days 60 and 65), are also apparent here, i.e. when considering only the component of change along the DSC.

**Extended Data Figure 10.**
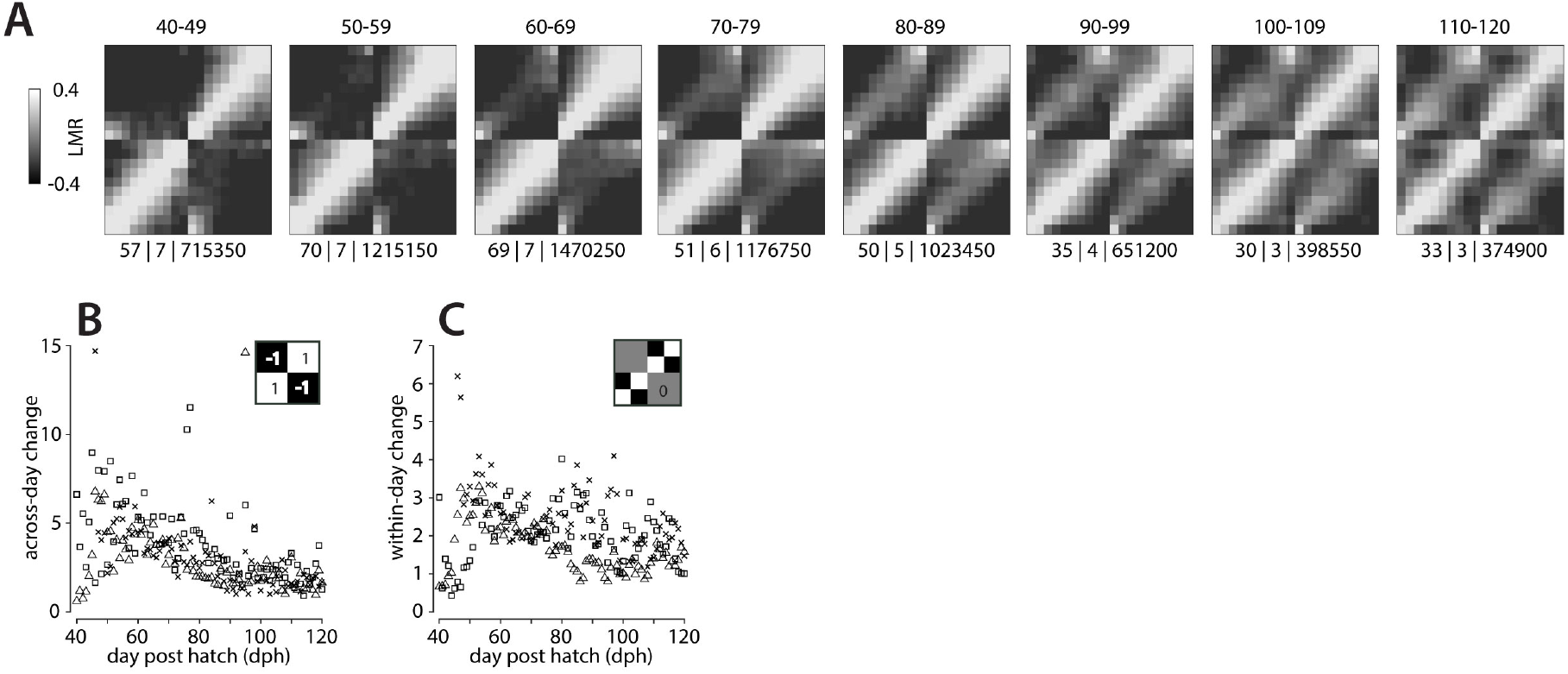
Two-day mixing matrices without stratification for 8 developmental phases. (**A**) Day-to-day mixing matrices for 8 phases during development. We binned renditions of each day into 10 consecutive periods based on production time (as in Fig. 3E), computed neighborhood mixing between all bins from two adjacent days, and averaged across day-pairs within a given developmental phase and birds. Numbers above each matrix indicate the range of days in each developmental period (start to end day post hatch, dph). Numbers below each matrix (separated by vertical bars) correspond to the number of mixing matrices that were averaged, the number of birds, and the total number of renditions across birds. Each 20-by-20 mixing matrix is analogous to one of the 10-by-10 squares along the diagonal of the stratified mixing matrices in Fig. 4A (e.g. model 1), but here is computed with finer time bins (10 periods vs. 5) and without separating renditions into strata. (**B**) Time-course of across-day change. Across-day change is obtained by multiplying a mask (inset) with measured day-to-day mixing-matrices (A) for a single pair of days and adding the entries of the resulting matrix (see Methods). Data from 3 animals with recordings up to dph 120 (symbols). (**C**) Timecourse of within-day change analogous to (B) but computed with a different mask (inset).

